# An autoregulatory feedback loop converging on H2A ubiquitination drives synovial sarcoma

**DOI:** 10.1101/2022.07.18.499263

**Authors:** Nezha S. Benabdallah, Vineet Dalal, Afroditi Sotiriou, R. Wilder Scott, Felix K.F. Kommoss, Anastasija Pejkovska, Ludmila Gaspar, Lena Wagner, Francisco J. Sánchez-Rivera, Monica Ta, Shelby Thornton, Torsten O. Nielsen, T. Michael Underhill, Ana Banito

## Abstract

The SS18-SSX fusion drives oncogenic transformation in synovial sarcoma by bridging SS18, a member of mSWI/SNF complex, to Polycomb repressive complex 1 (PRC1) target genes. Here we show that the SSX C-terminus, via its SSXRD domain, directs SS18-SSX chromatin binding independently of SS18. SSXRD specific targeting is mediated by interaction with mono ubiquitinated H2A (H2AK119ub1) and histone MacroH2A with which the fusion overlaps genome wide. Variant Polycomb Repressive Complex 1.1 (PRC1.1) acts as the main depositor of H2AK119ub1 and is therefore required for SS18-SSX occupancy. Importantly, the SSX C-terminus not only depends on H2AK119ub1 for localization but also further increases it by promoting PRC1.1 complex stability. Consequently, high H2AK119ub1 levels are a feature of murine and human synovial sarcomas. These results reveal an SSX/PRC1 autoregulatory feedback loop that reinforces fusion chromatin binding and therefore its oncogenic activity, and could play a role in a wider range of cancers and physiological settings where SSX proteins are overexpressed.

## INTRODUCTION

The Polycomb group (PcG) repressive system is an indispensable regulator of precise gene expression in all eukaryotes. Two classes of Polycomb proteins form distinct multi-protein complexes: Polycomb Repressive Complex 1 (PRC1), which mono-ubiquitylates histone H2A at lysine 119 (H2AK119ub1), and Polycomb Repressive Complex 2 (PRC2), which mono-, di-, and tri-methylates histone H3 at lysine 27 (H3K27me1, H3K27me2, and H3K27me3)^1,2^. Polycomb mediated regulation of gene expression is critical for embryogenesis as alterations in H2AK119ub1 and H3K27me3 levels lead to inappropriate expression of Polycomb target genes and embryonic defects^3–7^. The catalytic core of PRC1 is formed by RING1A or RING1B and one of six PCGF proteins, giving rise to distinct protein assemblies that comprise canonical PRC1 (cPRC1) or variant PRC1 (vPRC1) complexes^8,9^. vPRC1 complexes containing PCGF1/3/5/6 synergize to deposit the majority of the highly dynamic H2AK119ub1 mark^10–12^. Via H2AK119ub1 deposition, these complexes contribute globally to Polycomb domain formation through subsequent recruitment of PRC2 and H3K27me3. Ultimately, this triggers further recruitment of cPRC1 containing PCGF2/4 and CBX proteins, consolidating Polycomb-mediated gene repression^7,13–15^. PCGF1-containing complexes (PRC1.1) recognize unmethylated CpG islands via the ZF-CxxC domain of its KDM2B sub-unit^8,16–20^ and are responsible for H2AK119ub1 deposition at key developmental genes^17,21,22^. We recently showed that KDM2B function is hijacked in synovial sarcoma^23^ and several studies uncovered recurrent mutations in the PRC1.1 subunit BCOR in paediatric solid tumours, altogether suggesting a prominent role for PRC1.1 in tumour formation^24–29^.

Sarcomas are a group of cancers arising in soft tissues or bone that disproportionately affect children and young adults. Like other paediatric cancers, many types of sarcoma display a low mutational burden and are driven by dominant fusion oncoproteins involving chromatin associated regulators and transcription factors^30^. Synovial sarcoma, one of the more common soft tissue tumours in this class, is driven by the in-frame fusion of the mammalian SWI/SNF (mSWI/SNF or BAF) chromatin remodelling complex subunit SS18, where the last eight amino acids are replaced by the C-terminal 78 amino acids of SSX1, SSX2, or, rarely, SSX4^31,32^. Biochemical and proteomic studies have identified that SS18–SSX integrates into the mSWI/SNF via SS18 leading to the eviction of the tumour suppressor subunit SMARCB1 (also called BAF47 or hSNF5)^33,34^. This led to the idea that an altered mSWI/SNF complex is required for tumour formation^33,35^. However, later studies showed that SMARCB1 loss is not required for SS18-SSX driven tumourigenesis and rather it generates tumours with epithelioid sarcoma features in mice^35,36^. Thus, it is unclear to what extent mSWI/SNF complex deregulation is the defining event for synovial sarcoma identity.

We previously showed that SS18-SSX1 co-occupies KDM2B/PRC1.1 target sites and that KDM2B suppression disrupts SS18-SSX chromatin occupancy triggering proliferative arrest and acquisition of a fibroblast-like morphology^23^. Consequently, SS18-SSX guides the mSWI/SNF complex to Polycomb target genes leading to their aberrant activation^23,36^. Our previous work pinpointed PRC1.1 de-regulation as critical to sustain synovial sarcoma transformation. However, whether SS18-SSX recruitment onto chromatin at KDM2B sites is mediated by direct interactions with PRC1.1 members or by an indirect mechanism remained unclear.

Here we combine CRISPR/Cas9 gene-tiling screens, proteomics, FRET-based proximity assays and other molecular analyses to dissect SS18-SSX’s chromatin localizing mechanism. We show that the SSX C-terminus is solely responsible for SS18-SSX chromatin binding at its specific targets by interaction with H2AK119ub1 and histones MacroH2A. In addition to H2AK119ub1 deposition, we show that variant PRC1.1 can recruit histones MacroH2A, which to date lack a characterised chaperone, and is therefore critical to establish SS18-SSX chromatin localisation and maintenance. Furthermore, we uncover an autoregulatory feedback loop in which SS18-SSX both binds to and promotes H2AK119ub1 via two pathways - by up-regulating the expression of PRC1.1 members *BCOR* and *RYBP*, and by stabilising the PRC1.1 complex presence on chromatin via its SSX-C domain. This autoregulatory mechanism allow SS18-SSX to reinforce its own activity and results in acquisition of high H2AK119ub1 levels during synovial sarcoma tumourigenesis.

## RESULTS

### The SSX C-terminus is sufficient for tight and specific SS18-SSX chromatin binding

To map protein domains in SS18-SSX that are essential for tumour maintenance, we performed a CRISPR-Cas9 knockout screen using a gene-tiling sgRNA library covering the entire SS18 and SSX1 coding sequences **(Fig. 1a)**. In this assay, sgRNAs targeting DNA sequences coding for essential protein domains often result in a more significant dropout, since even small inframe indels in these regions are likely to affect protein function and cell fitness^37,38^. We screened for critical SS18-SSX1 domains in the HS-SY-II synovial sarcoma cell line and used ProTiler to map CRISPR knockout hyper-sensitive (CKHS) regions^39^. As expected, sgRNAs targeting SS18 were generally depleted with the exception of those targeting a region that is not present in SS18’s shorter isoform (aa 295-325). Still, a clear CKHS region was identified at the SSX C-terminus corresponding to the highly conserved SSXRD (SSX Repression Domain)^40,41^ **(Fig. 1b, Extended Fig.1a,b)**. These results suggest that this region, consisting in the last 34 amino acids of SS18-SSX1, is most critical for its oncogenic function.

**Figure 1:**
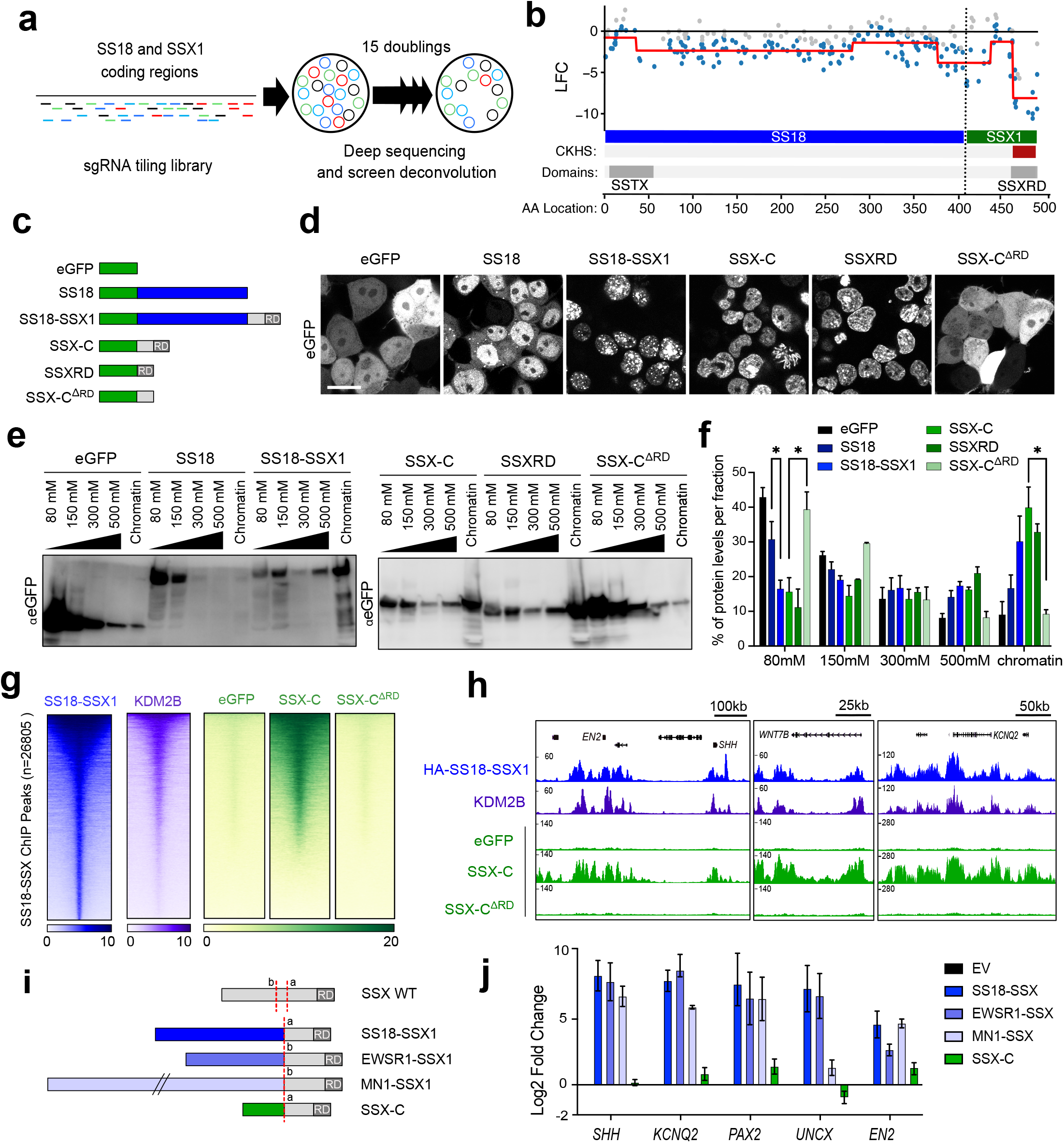
SSX C-terminus directs tight and specific SS18-SSX chromatin binding. **a**) Layout of CRISPR-Cas9 knockout gene-tilling screen. **b)** Mapping of CRISPR knockout hyper-sensitive (CKHS) regions in SS18-SSX1 using ProTiler based on Iog2 fold changes (LFC) of sgRNAs representation in HS-SY-II synovial sarcoma cells. The CKHS region is highlighted in dark red and corresponds to the SSXRD PFAM sequence (PF09514). **c)** Schematic representation of eGFP (green) fused constructs for SS18, SS18-SSX1, SSX-C (78aa of SSX1 present in the SS18-SSX1 fusion), SSXRD (last 34aa of SSX-C) or SSX-CΔRD (SSX-C with a deletion of the SSXRD). **d)** Live confocal imaging of the eGFP-fused constructs in HEK193T cells. Scale bar corresponds to 20μm. **e)** Salt extraction assay in HEK293T expressing the various eGFP constructs. The proteins are detected using an eGFP antibody. **f)** Percentage of total protein levels per fraction in two or three biological replicates. Data represents the mean and the standard error of the mean (S.E.M). Asterisks represent p-values of paired one-tailed t-test between groups, * p< 0.05. **g)** Heatmaps for HA-SS18-SSX1, KDM2B ChlP-seq from Banito et al., 2018 and eGFP ChIP in HS-SY-II cells expressing eGFP fused SSX-C or SSX-C ^ΔRD^. Heatmaps represent ChlP-seq signals over HA-SS18-SSX1 broad peaks (n=26805). Rows correspond to ±5-kb regions across the midpoint of each HA-enriched region, ranked by increasing signal in HS-SY-II cells. **h)** Gene tracks for HA-SS18-SSX1, KDM2B and eGFP ChlP-seq at the *EN2, WNT7B* and *KNCQ2* loci. **i)** Schematic representing new synovial sarcoma fusions. SS18-SSX1 and the SSX-C contain the canonical breakpoint “a”, while EWSR1-SSX1 and MN1-SSX1 exhibit an alternative breakpoint “b”. **j)** qRT-PCR displaying Log2 fold change of mRNA levels relative to *GAPDH* in mesenchymal stem cells (MSCs) expressing the new fusion constructs and controls. Values are normalized to empty vector expression in three biological replicates. Data represents the mean and the standard error of the mean (S.E.M).

To explore the role of SSXRD, we generated constructs containing eGFP fused to the SSX1 C-terminal region present in SS18-SSX1, with or without a SSXRD deletion (SSX-C^ΔRD^ and SSX-C, respectively) or the SSXRD alone. SS18 and SS18-SSX1 eGFP fusions were used as controls **(Fig. 1c)**. When expressed in HEK293T, eGFP-SS18 exhibits both nuclear and cytoplasmic localisation. In contrast, eGFP-SS18-SSX1, SSX-C and SSXRD are exclusively detected in the nucleus. Most importantly, SSX-C nuclear localisation depends on the SSXRD domain, as eGFP-SSX-C^ΔRD^ loses the exclusive nuclear pattern. This further supports the presence of a nuclear localisation signal in the SSXRD **(Fig. 1d)**^42^ and is suggestive of a key role of this region in mediating chromatin interaction.

To assess the presence and strength of the association between SSX1 C-terminus and chromatin we performed sequential salt extractions. Chromatin-associated proteins become more soluble with increasing concentrations of NaCl, with proteins strongly bound to DNA, such as histones H2A/H2B, eluting at high salt concentrations (>800 mM)^43–45^. We observed that SSXRD is tightly bound to chromatin and is resistant to 500mM salt extraction. Similarly, SSXRD-containing GFP fusions, but not eGFP-SSX-C^ΔRD^, are predominantly present in the chromatin fraction indicating that SS18-SSX strongly binds chromatin via this domain at the SSX C-terminus **(Fig. 1e, f)**. To determine if SSX-C alone can reproduce the SS18-SSX-specific genome wide occupancy, we performed chromatin immunoprecipitation sequencing (ChIP-Seq) of eGFP-SSX-C overexpression in HS-SY-II synovial sarcoma cells. This revealed a clear overlap with previously identified SS18-SSX/KDM2B bound regions which was abolished in the absence of the SSXRD domain **(Fig. 1g, h)**. These results indicate that the SSX c-terminus, via its SSXRD, binds specific regions in the genome and determines SS18-SSX localisation independently of SS18 and therefore of the mSWI/SNF complex.

In line with these results, recent studies identified SSX fusions in synovial sarcoma involving alternative activators such as EWSR1 and MN1^46^. This potentially indicates that recruitment of other partners fused to the SSX C-terminus can result in the activation of PRC1.1 target genes. To test this hypothesis, we expressed *EWSR1-SSX1* and *MN1-SSX1* in human mesenchymal stem cells **(Fig. 1i)**. Similar to SS18-SSX1, expression of the new fusions resulted in specific up-regulation of PRC1.1 target genes that are part of the synovial sarcoma gene signature **(Fig. 1j)**^23^. These results demonstrates that induction of Polycomb target genes can be achieved by recruitment of other transcriptional activators, and that the SSXRD is the key element that defines a synovial sarcoma signature.

### The SSXRD acidic tail links SSX to histone H2AK119ub1 and MacroH2A domains

Consistent with a tight SSXRD-dependent chromatin binding, SS18-SSX and SSX-C remain associated with condensed chromosomes during mitosis **(Extended Fig. 1c)**^47^. However, neither KDM2B, PCGF1, nor RING1B binds the mitotic chromosome to the same extent as SSXRD, suggesting that the SSX C-terminus can bind chromatin independently of the presence of these PRC1.1 components **(Extended Fig. 1d)**. To identify candidate factors mediating SSX-C/SSXRD chromatin binding, we investigated the common interactome of SSX-C and SSXRD by performing eGFP co-immunoprecipitation followed by mass spectrometry **(Fig. 2a)**. Enriched proteins were defined by normalization to SSX-C^ΔRD^ pull-down which, as expected, showed a strong overlap with the eGFP-only negative control. Noticeably, histones were highly represented in both SSX-C and SSXRD top interactors, with higher enrichment than PRC1 or PRC2 components, indicating that SSX-C can bind chromatin via a direct histone interaction **(Fig. 2b)**.

**Figure 2:**
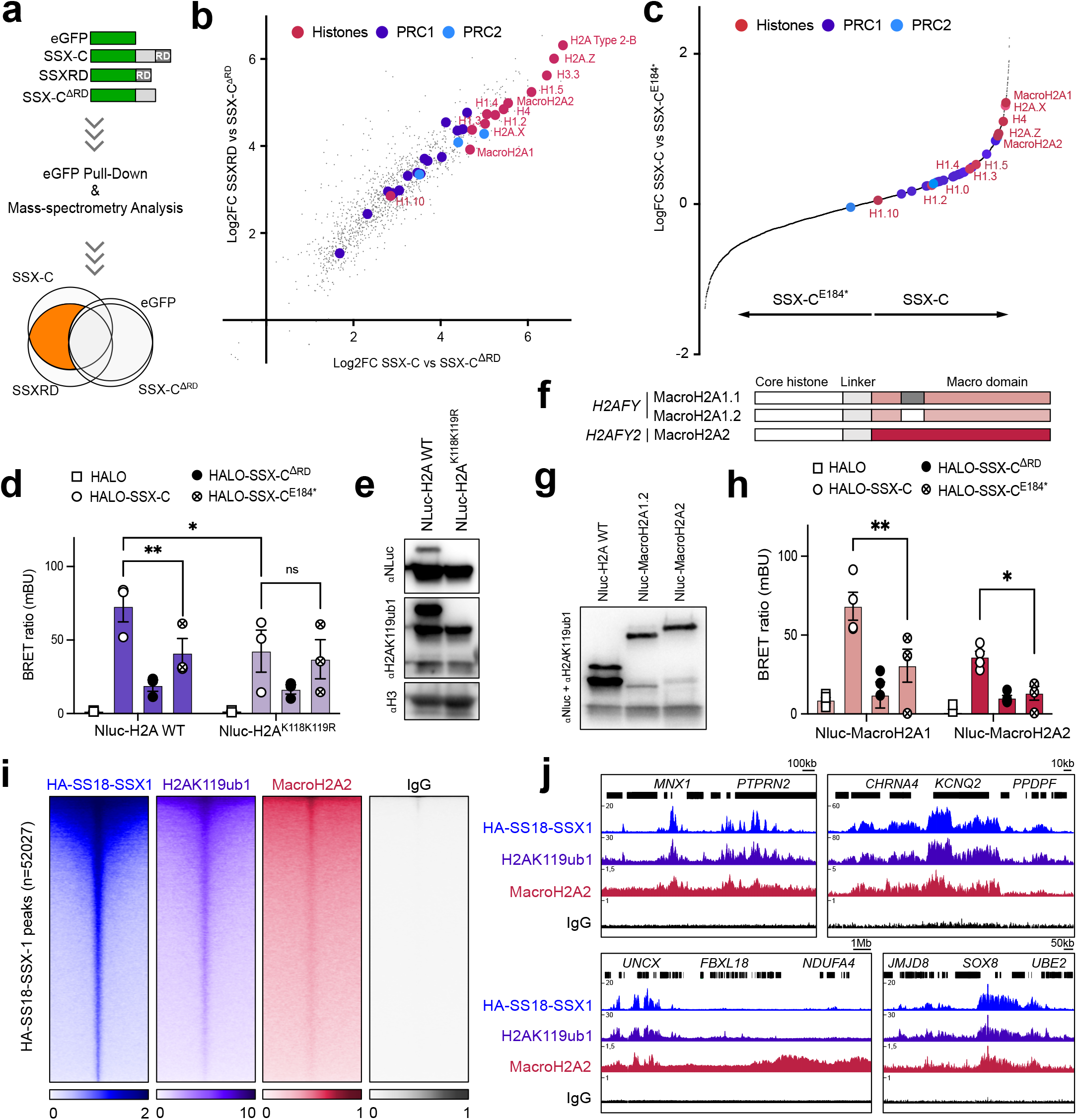
The SSXRD acidic tail links SSX to histone H2AK119ub1 and MacroH2A domains. **a)** Schematic representing the layout of the eGFP pull down and mass-spectrometry analysis. Common SSX-C and SSXRD enriched hits (highlighted in orange) were identified. **b)** Log2 fold change correlation plot of eGFP-SSXRD and eGFP-SSX-C mass spectrometry data following eGFP pull down in HS-SY-II cells. Data was normalized to eGFP-SSX-C ^ΔRD^. **c)** Log fold change plot between eGFP-SSX-C and eGFP-SSX-C^E184^* mass spectrometry data following eGFP pull down in two biological replicates. Data was normalized to eGFP. **d)** BRET ratio (mBU) in Nluc-H2A or Nluc-H2A^K118K119R^ transfected HEK293T cells expressing empty vector HALO, HALO-SSX-C, HALO-SSX-C^ΔRD^ or HALO-SSX-C^E184^*. Values represent 3 biological replicates. Asterisks represent p-values of paired one-tailed t-test between groups ** p< 0.01, * p< 0.05. **e)** Western blot of histone acid extracts from HEK293T cells transfected with either Nluc-H2A or Nluc-H2A^K118K119R^ revealed with NLuc, H2AK119ub1 and H3 antibodies. **f)** Illustration of the two MacroH2A genes, *H2AFY* encoding the two isoforms MacroH2A1.1 and MacroH2A1.2 which differ by one exon (grey/white box) within the Macro domain (pink) and *H2AFY2* encoding MacroH2A2. **g)** Western blot of histone acid extracts from HEK293T cells transfected with either Nluc-H2A, Nluc-MacroH2A1.2 or Nluc-MacroH2A2. Detection was performed using NLuc mixed with H2AK119ub1 and H3 antibodies. **h)** BRET ratio (mBU) in Nluc-H2A, Nluc-macroH2A1.2 or Nluc-macroH2A2 transfected HEK293T cells expressing empty vector HALO, HALO-SSX-C, HALO-SSX-C^ΔRD^ or HALO-SSX-C^E184^*. Values represent 4 biological replicates. Asterisks represent p-values of paired one-tailed t-test between groups (* p< 0.05, ** p< 0.01). **i)** Heatmaps of HA-SS18-SSX1, H2AK119ub1 and MacroH2A2 CUT&RUN signals over HA-SS18-SSX1 peaks in HS-SY-II cells (n=52027). Rows correspond to ±5-kb regions across the midpoint of each signal, ranked by increasing signal. **j)** Gene tracks for HA-SS18-SSX1, H2AK119ub1 and MacroH2A2 CUT&RUN in HS-SY-II cells at the *MNX1, KCNQ2, UNCX* and *SOX8*.

Recent *in vitro* biochemistry studies showed the ability of SSX-C to bind the nucleosome acidic patch, with a preference for H2AK119ub1-modified nucleosomes conferred by the last 5 amino acids (EEDDE) of the SSXRD^41,48^. We generated an eGFP-fused SSX-C mutant lacking the last 5 amino acids of SSXRD (SSX-C^E184*^). Accordingly, when expressed in HEK293T cells which contain several copies of the X chromosome, SSX-C^E184*^ loses the SSXRD specific co-localisation to H2AK119ub1-rich Barr bodies, whilst maintaining a nuclear localisation pattern **(Extended Fig. 2a)**. Therefore, to further dissect SSX-C interactors driving SSX specific chromatin occupancy, we compared eGFP pull-downs of SSX-C with that of SSX-C^E184*^. Remarkably, our proteomic analysis indicated that in addition to histone H2A, other histones H2A variants such as MacroH2A1, H2AX, H2AZ and MacroH2A2, appeared to be strong interactors **(Fig. 2c)**.

To investigate the H2AK119ub1/SSX-C interaction in live cells, we performed NanoBret, a protein-protein interaction assay based on bioluminescence resonance energy transfer (BRET)^49,50^. We detected an interaction of the SSX-C (Halo-SSX-C) when co-expressed with histone H2A fused to Nano Luciferase (NLuc-H2A). This interaction was dependent on the SSXRD domain and was diminished in the SSX-C^E184*^ mutant **(Fig. 2d)**. These results support a role for the EEDDE-containing SSXRD domain in targeting of SS18-SSX to nucleosomes. Next, to test if H2AK119ub1 plays a role in SSXRD chromatin targeting in living cells we performed NanoBret, this time expressing either wild-type histone H2A (NLuc-H2A) or a mutant H2A that cannot be ubiquitinated, (NLuc-H2A^K118K119R^)^51^ **(Fig. 2e)**. The absence of ubiquitination decreased the ability of SSX-C to interact with the nucleosome *in vivo* indicating that H2AK119ub1 plays an active role in specifying SS18-SSX chromatin occupancy **(Fig. 2d)**.

As MacroH2A histones have been previously linked with Polycomb co-occupancy, transcriptional repression and X chromosome inactivation^52–56^ we set out to test this additional link between PRC1 and SS18-SSX activity in synovial sarcoma. MacroH2A histones contain a homologous histone domain that is integrated in the nucleosome, followed by a linker region and a large macro domain that extends outside of the nucleosome^57^. MacroH2A1 and MacroH2A2 proteins are encoded by *H2AFY* and *H2AFY2* with MacroH2A1 presenting with two isoforms (MacroH2A1.1 and MacroH2A1.2) that differ in a single exon **(Fig. 2f)**. We first confirmed the co-localisation of the three different MacroH2A histones, MacroH2A1.1, MacroH2A1.2 and MacroH2A2, with SSX-C at the inactive X chromosome in HEK293T cells **(Extended Fig. 2b)**. Next, we performed NanoBret using NLuc-MacroH2A1.2 or Nluc-MacroH2A2 and we observed an interaction between MacroH2A1.2/MacroH2A2 and SSX-C that was significantly diminished in SSX-C^E184*^ **(Fig. 2g, h)**.

To gain insight into global chromatin co-occupancy of SS18-SSX1, H2AK119ub1 and MacroH2A2, we performed CUT&RUN^58^ using a HS-SY-II synovial sarcoma cell line where SS18-SSX1 is endogenously labelled with an HA-tag^23^. This revealed that MacroH2A2 is present at sites highly enriched in H2AK119ub1 and HA-SS18-SSX1 (*MNX1, KCNQ2, UNCX, SOX8)* **(Fig. 2i, j)**. However, characteristic MacroH2A2 broad domains^56,59^ did not correlate with SS18-SSX, nor H2AK119ub1 **(Fig. 2j)** indicating that MacroH2A2 alone is not sufficient to cause SS18-SSX targeting. Accordingly, SS18-SSX1 and H2AK119ub1 genome occupancy profiles strongly correlate, whilst MacroH2A2 does so moderately **(Extended Fig. 2c)**. Together, our results highlight that a very specific chromatin environment containing both H2AK119ub1-modified nucleosomes and histone variant MacroH2A underlies SSX C-terminus chromatin binding.

### PRC1.1 deposits H2AK119ub1 and MacroH2A and mediates SS18-SSX recruitment independently of PRC2

The chromatin environments bound by SSXRD that are rich in H2AK119ub1 and MacroH2A are also co-occupied by several other chromatin regulators. vPRC1 deposits H2AK119ub1 which is recognized and bound by PRC2. This leads to H3K27me3 deposition which in turn results in cPRC1 recruitment^7,13–15^.

To dissect the hierarchy of SS18-SSX targeting at Polycomb sites, we first assessed whether KDM2B, which mediates recruitment of PRC1.1, is sufficient to recruit SS18-SSX onto chromatin. To this end, we took advantage of a previously described artificial targeting approach where KDM2B is fused to the methyl binding domain (MBD) of MBD1 leading to its re-targeting to regions of densely methylated DNA such as pericentromeric heterochromatin **(Fig. 3a)**^60,61^. Additionally, a critical residue in the zf-CXXC DNA binding domain of KDM2B^62^ is mutated so that the MBD-fused protein can only bind methylated DNA (MBD-KDM2B^K643A^ referred to as MBD-KDM2B). MBD fused to Luciferase (MBD-Luc) was used as control to assess specific targeting **(Fig. 3a)**. We first confirmed the correct tethering of the MBD-fused proteins to heterochromatin using immunofluorescence in the human synovial sarcoma cell line HS-SY-II harbouring endogenously HA tagged SS18-SSX^23^. We observed a specific co-localisation of the MBD constructs, marked by a V5 tag, to heterochromatin protein 1 (HP1) foci **(Extended Fig. 3a)**. As expected, MBD-KDM2B, but not MBD-Luc, was able to recruit PRC1.1 components BCOR and PCGF1, resulting in H2AK119ub1 deposition at V5/HP1 foci **(Fig. 3b, Extended Fig. 3b)**. Most importantly, MBD-KDM2B was sufficient to recruit SS18-SSX1 **(Fig. 3c)**. Interestingly, MBD-KDM2B did not lead to accumulation of H3K27me3 at the V5/HP1 foci **(Fig. 3d)**, indicating that PRC2 could not be involved in SS18-SSX recruitment. To further dissect the role of PRC1.1 and PRC2 in SS18-SSX recruitment, we employed genetic knock-out of the PRC1.1 or PRC2 components using CRISPR/Cas9-directed mutagenesis^38,63,64^ **(Extended Fig. 3c)**. Remarkably, depleting the PRC1.1 components BCOR or PCGF1 but not the PRC2 components EED or EZH2 significantly reduced SS18-SSX1 recruitment and H2AK119ub1 deposition mediated by MBD-KDM2B **(Figure 3e, Extended Fig. 3d)**. Consistent with our observations, MBD-KDM2B tethering did not lead to recruitment of PRC2 components **(Extended Fig. 3e)**. Together these results indicate that SS18-SSX1 targeting can be initiated by KDM2B, relies on an intact PRC1.1 complex but is independent from PRC2 activity.

**Figure 3:**
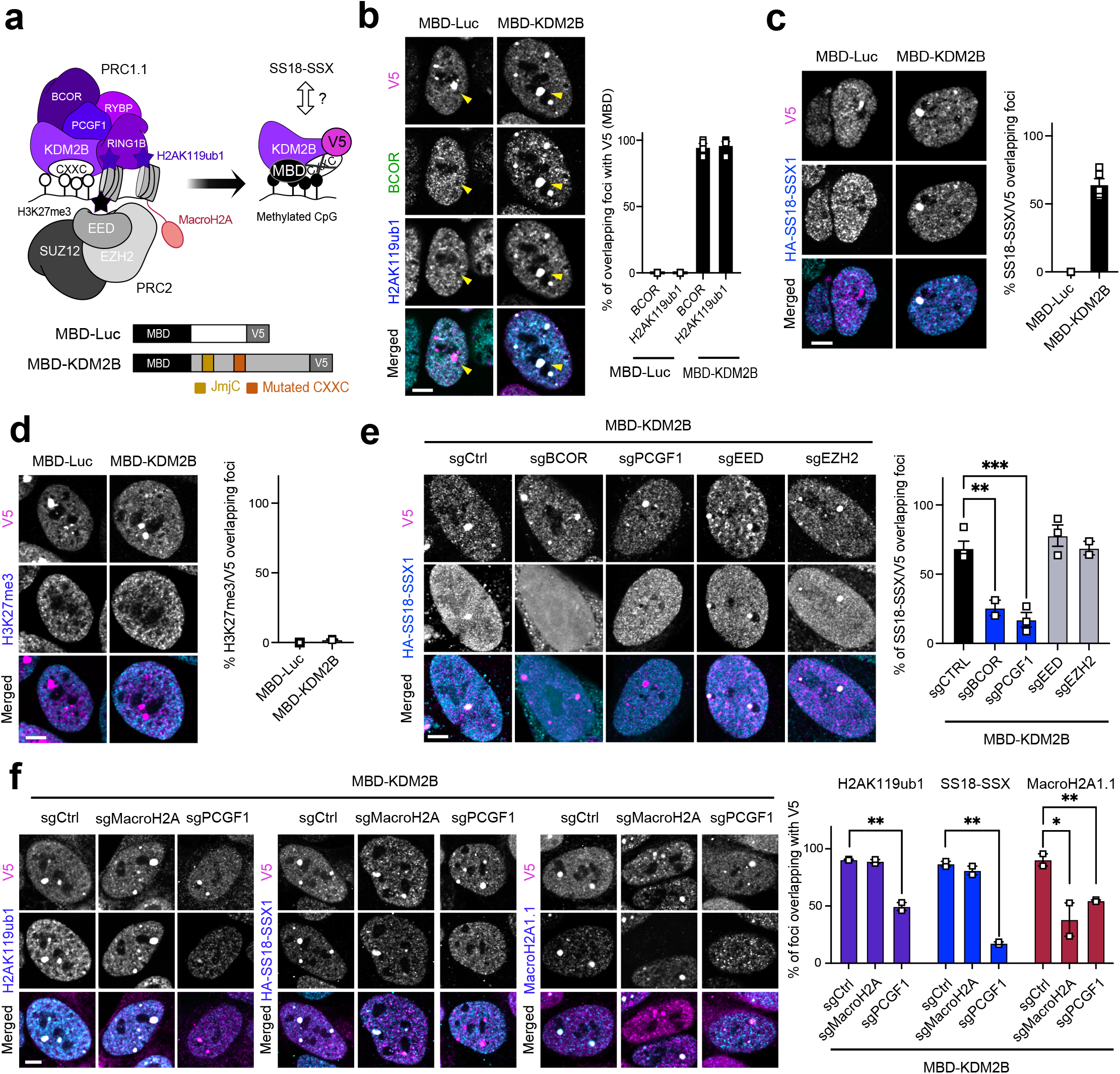
PRC1.1 deposits H2AK119ub1 and MacroH2A and regulates SS18-SSX recruitment independently of PRC2. **a)** Up, Representation of the methyl binding domain (MBD)-mediated targeting approach of proteins to methylated CpG. Here, the CXXC mutated KDM2B is redirected to methylated CpG via the MBD. Bottom, Schematic representing the MBD (black square) fusion constructs for Luciferase control (Luc) and KDM2B. The KDM2B long isoform contains the histone demethylase JmjC domain (gold box) and a mutated CXXC domain (dark orange). Both constructs contain a V5 tag. **b)** Left, Immunofluorescence of human synovial sarcoma (HS-SY-II) cells displaying the MBD constructs (V5, magenta), BCOR (green) and H2AK119ub1 (cyan). Yellow arrow heads point to the MBD foci. Scale bars represents 5μm throughout the figure. Right, quantification of the percentage of BCOR or H2AK119ub1 foci overlapping a V5 foci in 3 or 4 biological replicates. Data represents the mean and the standard error of the mean (S.E.M). **c)** Left, Immunofluorescence for V5 (magenta) and SS18-SSX1 (HA, cyan). Right, quantification of the percentage of HA (SS18-SSX1) foci overlapping a V5 foci in 2 to 5 biological replicates. **d)** Left, Immunofluorescence for V5 (magenta) and H3K27me3 (cyan). Right, quantification of the percentage of BCOR or H3K27me3 foci overlapping with V5 foci in 2 biological replicates. Data represents the mean and S.E.M. **e)** Left, Immunofluorescence of MBD-KDM2B (V5, magenta) in the presence of different sgRNAs (eGFP background fluorescence) with SS18-SSX1 (HA, cyan) in in HS-SY-II-Cas9 cells. Right, quantification of the percentage of HA (SS18-SSX1) foci overlapping a V5 foci in 2 to 4 biological replicates. Data represents the mean and S.E.M. Asterisks represent p-values of unpaired one-tailed t-test between groups (** p< 0.01 and *** p< 0.001). **f)** Left, Immunofluorescence images of MBD-KDM2B (V5, magenta) with SS18-SSX or H2AK119ub1 or MacroH2A1.1 (cyan) in HS-SY-II-Cas9 cells expressing sgCtrl, sgMacroH2A (targeting both histone genes *H2AFY* and *H2AFY2)* or sgPCGFI. Right, quantification of the percentage of foci overlapping MBD-KDM2B foci in 2 biological replicates. Data represents the mean and S.E.M. Asterisks represent p-values of unpaired one-tailed t-test between groups (* p< 0.05 and ** p< 0.01).

MacroH2A histones deposition has been mainly associated with PRC2 even though the chaperone orchestrating its deposition at these sites remains unknown^65^. We took advantage of the MBD-KDM2B tethering assay to dissect the interplay between PRC1.1, MacroH2A and H2AK119ub1 deposition. We observed an unanticipated concomitant deposition of MacroH2A histones that in synovial sarcoma cells occurs independently of H3K27me3 **(Extended Fig. 3f)**, suggesting that PRC1.1 could additionally promote the SS18-SSX oncofusion recruitment by fostering MacroH2A incorporation.

To test their requirement for tumour cell maintenance, we removed both MacroH2A isoforms (sg*H2AFY* combined with sg*H2AFY2*, referred to as sgMacroH2A) or *PCGF1* in HS-SY-II synovial sarcoma cells and in KHOS-240s osteosarcoma cells using CRISPR/Cas9-directed mutagenesis with sgRNAs carrying an eGFP reporter. We observed that, like sgPCGF1, sgMacroH2A specifically impacted synovial sarcoma cell growth, albeit with a much milder effect in the case of MacroH2A **(Extended Fig. 3g)**. We then assessed whether MacroH2A is required for *de novo* recruitment of SS18-SSX1 using the MBD assay, we observed that only PCGF1 knockout but not MacroH2A removal led to a significant decrease in SS18-SSX recruitment **(Fig. 3f)**. Importantly, PRC1.1 removal affected the accumulation of both H2AK119ub1 and MacroH2A, indicating that MacroH2A deposition depends on PRC1.1 activity in synovial sarcoma cells **(Fig. 3f, g, Extended Fig. 3h)**. We therefore uncovered a role for PRC1.1 in promoting accumulation of MacroH2A histones independently of PRC2. Thus, PRC1.1 is sufficient to establish H2AK119ub1 and MacroH2A-rich chromatin environments that are recognized by SSXRD.

### PRC1.1 controls global H2AK119ub1 deposition and is required for SS18-SSX occupancy

As several PRC1 complexes can mediate H2AK119ub1 deposition, it raises the question of whether PRC1.1 inhibition alone in synovial sarcoma cells is able to deplete the mark resulting in loss of SS18-SSX at its target sites. To assess the global effect of PCGF1 removal on SS18-SSX recruitment and H2AK119ub1 genome-wide, we used CUT&RUN to profile PCGF1 knockout in HS-SY-II and SYO-1 cells, harbouring SS18-SSX1 and SS18-SSX2 fusions respectively. Strikingly, PCGF1 knockout led to a global decrease in H2AK119ub1 deposition alongside a strong reduction in SS18-SSX1/2 chromatin occupancy in both cell lines illustrating the pivotal role of variant PRC1.1 in SS18-SSX chromatin maintenance **(Fig. 4a, b, c)**.

**Figure 4:**
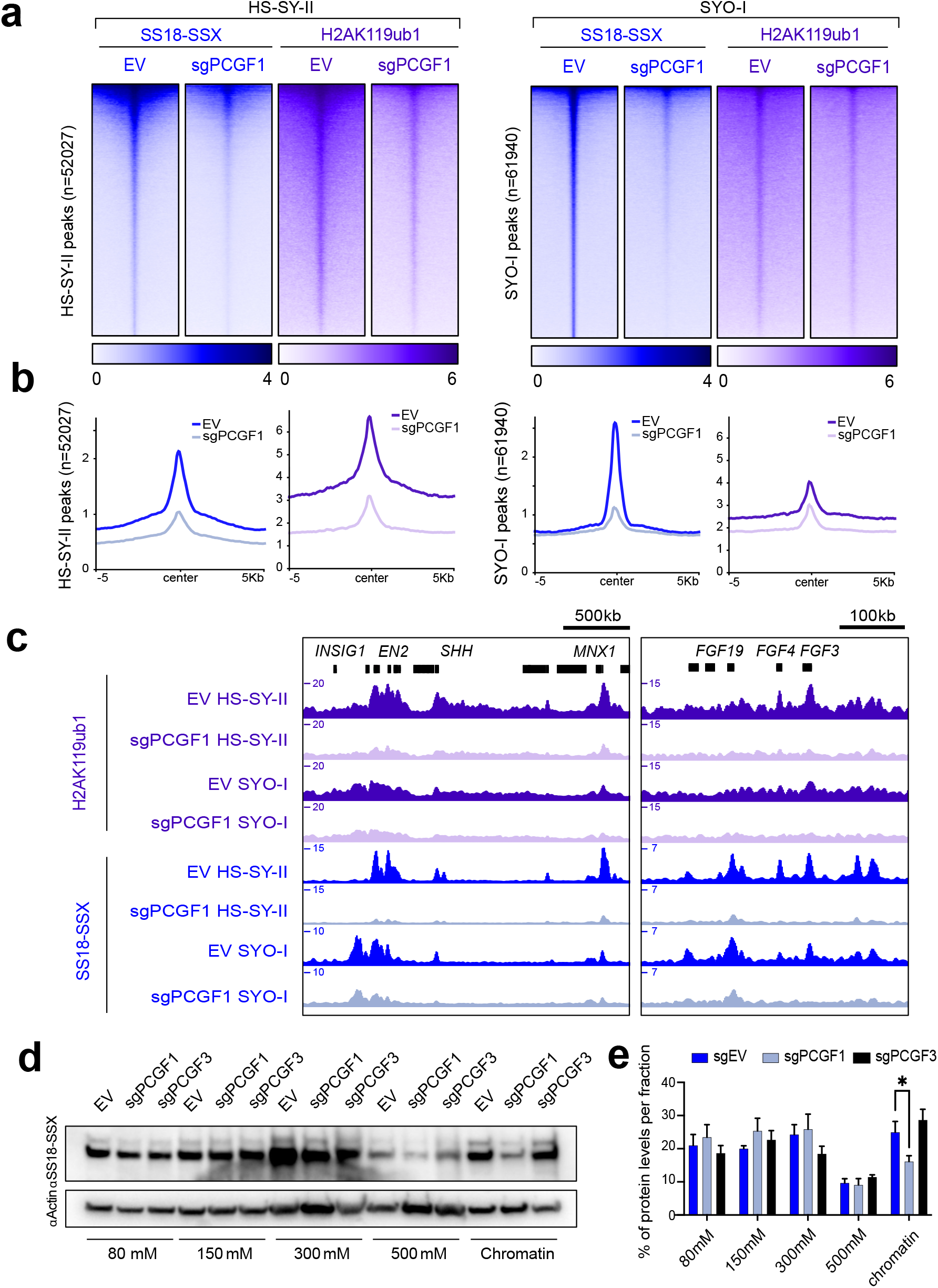
PRC1.1 controls global H2AK119ub1 deposition and regulates SS18-SSX recruitment. **a)** Heatmaps of H2AK119ub1 (purple) or SS18-SSX (blue) CUT&RUN signals in HS-SY-II (left) and SYO-I (right) Cas9 cells expressing empty sgRNA as control (EV) or targeting PCGF1 (sgPCGFI). Both heatmaps represent CUT&RUN signals over HA-SS18-SSX1 peaks in HS-SY-II (left, n=52027) or SS18-SSX2 peaks in SYO-I (right, n=61940). Rows correspond to ±5-kb regions across the midpoint of each enriched region, ranked by increasing signal. **b)** H2AK119ub1 and SS18-SSX CUT&RUN score distributions across HA-SS18-SSX1 peaks in HS-SY-II (left, n=52027) or the SS18-SSX2 peaks in SYO-I (right, n=61940). **c)** Gene track for H2AK119ub1 and SS18-SSX CUT&RUN at the *EN2-SHH-NOM1* and *FGF4-FGF3* loci. **d)** Salt extraction assay displaying SS18-SSX1 levels by western blot in HS-SY-II-Cas9 cells expressing an empty vector (EV) or sgRNAs against PCGF1 or PCGF3. **e)** Quantification of the SS18-SSX protein distribution in the various salt extraction fractions. Data represents the mean and S.E.M for the percentage of total protein per fraction in 3 biological replicates. Asterisks represent p-values of paired one-tailed t-test between groups (* p< 0.05).

We next assessed if PCGF3, a member of another PRC1 variant, PRC1.3, and a known dependency in synovial sarcoma (https://depmap.org/portal/), can impact SS18-SSX chromatin binding to the same extent as PCGF1. To compare the global action of PCGF1 versus PCGF3 in an unbiased manner, we performed chromatin salt extraction on SS18-SSX in both HS-SY-II and SYO-I synovial sarcoma cell lines following either PCGF1 or PCGF3 knockout. Whereas removal of PCGF1 abrogated SS18-SSX presence on chromatin, PCGF3 removal had no effect. Hence, although PCGF3 is required for synovial sarcoma maintenance, our results indicate that it does not affect SS18-SSX global chromatin binding suggesting instead an alternative role for PRC1.3 in this context **(Fig. 4 d, e)**. Our data show that PRC1.1 acts as the main depositor of H2AK119ub1 and is therefore needed for SS18-SSX chromatin binding.

### SS18-SSX enforces H2AK119ub levels by promoting expression of PRC1.1 components

Interestingly, *BCOR*, a PRC1.1 subunit that regulates complex assembly and activity^66,67^, is upregulated in synovial sarcoma tumour samples^68,69^, suggesting an interplay between SS18-SSX and PRC1.1. To investigate this, we looked into published RNA sequencing data where SS18-SSX1 expression is induced in naïve human fibroblasts (CRL7250)^36^. There, SS18-SSX1 expression resulted in increased *BCOR* and *RYBP* mRNA levels **(Fig. 5a)**. Reciprocally, SS18-SSX knock-down in the two synovial sarcoma cell lines HS-SY-II (SS18-SSX1) and SYO-I (SS18-SSX2) showed a concomitant decrease in *BCOR* and, to less extent, *RYBP* expression levels^36^ **(Fig. 5b)**. Similarly, when we expressed SS18-SSX1 in mesenchymal stem cells or removed SS18-SSX1 in synovial sarcoma cells, we observed increased or reduced BCOR and PCGF1 protein levels, respectively **(Fig. 5c, d)**. This suggests that SS18-SSX directly regulates *BCOR* and *RYBP* transcription levels, and indeed both genes are direct targets of the oncofusion protein **(Fig. 5e)**.

**Figure 5:**
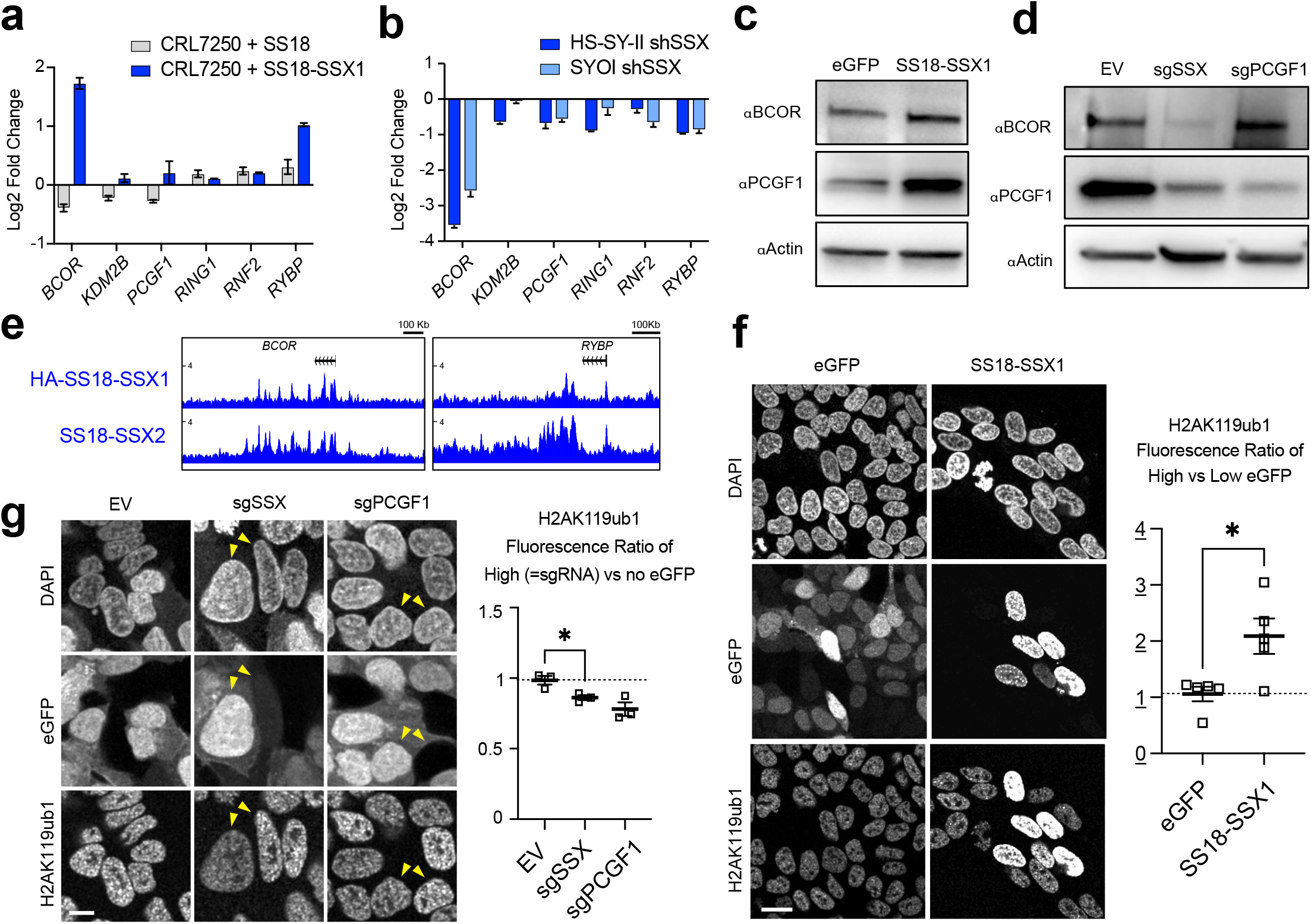
SS18-SSX regulates expression of PRC1.1 components and increases H2AK119ub levels. **a)** Log2 Fold change of RPKM values from RNA sequencing in CRL7250 cells expressing SS18 or SS18-SSX1 compared to naive cells in two biological replicates. Data from (McBride et al., 2018). **b)** Log2 Fold change of RPKM values from RNA sequencing in HS-SY-II and SYO-I cells after knockdown of SS18-SSX compared to shCtrl cells in two biological replicates. Data from (McBride et al., 2018). **c, d)** Western blot of whole cell extracts from mesenchymal stem cells expressing eGFP or eGFP-SS18-SSX1 (c); and from HS-SY-II-Cas9 cells expressing empty sgRNA vector (EV) or sgRNA against SSX or PCGF1 (sgSSX, sgPCGFI) (d). Proteins were detected using BCOR and PCGF1 antibodies and Beta-actin was used as loading control. **e)** Gene tracks for HA-SS18-SSX1 and SS18-SSX2 CUT&RUN at the *BCOR* and *RYBP* loci. **f)** Left, Immunofluorescence of eGFP-fused constructs (eGFP, eGFP-SS18-SSX1) expressed in HS-SY-II with eGFP signals (green) and nucleus stained with DAPI and H2AK119ub1. Right, quantification of H2AK119ub1 fluorescence ratio in high versus low eGFP cells in 5 biological replicates. Data represents the mean and the S.E.M. Asterisks represent p-values of paired one-tailed t-test between groups (* p< 0.05). g) Left, H2AK119ub1 immunofluorescence in HS-SY-II-Cas9 cells expressing empty sgRNA vector (EV) or sgRNA against SSX or PCGF1 (sgSSX, sgPCGFI). sgRNA expressing cells are positive for eGFP. Cells were mixed with non sgRNA expressing cells for direct comparison of H2AK119ub1 signal intensity (yellow arrow heads). Right, quantification of the H2AK119ub1 fluorescence ratio in eGFP (=sgRNA) versus no eGFP cells in 3 biological replicates. Data represents the mean and the standard error of the mean (S.E.M). Asterisks represent p-values of paired one-tailed t-test between groups (* p< 0.05). Scale bars represents 10μm. Scale bars represents 20μm.

Consistent with a role for the BCOR and RYBP subunits in supporting PRC1.1 assembly and activity^66,67,70^, overexpression of the eGFP-SS18-SSX constructs in synovial sarcoma cells leads to higher H2AK119ub1 levels in a manner that correlates with the eGFP-fusion expression levels **(Fig. 5f)**. Of note, the same is not observed when expressing an eGFP-only control, where the H2AK119ub1 staining remains homogeneous regardless of the amount of construct in the cell. This indicates that SS18-SSX-mediated induction of PRC1.1 components leads to a detectable increase in H2AK119ub1. Reciprocally, by removing SS18-SSX in synovial sarcoma cells using an sgRNA against SSX, we also observed a reduction of H2AK119ub1 levels **(Fig. 5g)**. Importantly, while SS18-SSX inhibition impacts PCGF1 protein levels, it does not regulate its transcription **(Fig. 5a, b)**. These results indicate that SS18-SSX controls expression of the PRC1.1 member *BCOR* and is also able to promote PRC1.1 protein levels via an alternative mechanism.

### SSX-C increases PRC1.1 stability thus reinforcing H2AK119ub1 levels and SS18-SSX occupancy

Since SSX-C does not act as a transcriptional activator **(Fig. 1j)** but directly interacts with chromatin, we hypothesized that it could augment PRC1.1 protein levels by increasing stabilization of the complex on chromatin. First, we observed that overexpression of SSX-C alone is also able to induce BCOR levels in synovial sarcoma cells **(Fig. 6a)**. Similarly, SSX-C overexpression in mesenchymal stem cells, recapitulates the increase in BCOR levels **(Extended Fig. 6a)**. Of note, SSX-C overexpression does not induce *BCOR* transcription indicating that this increase occurs at the protein level **(Extended Fig. 6b)**. Sequential chromatin washes in HEK293T cells and chromatin salt extractions in HS-SY-II cells showed that SSX-C expression increases the presence of the PRC1.1 proteins BCOR and PCGF1 on the chromatin while decreasing their presence in more soluble fractions **(Fig. 6b, c and Extended Fig 6c)**. Such results suggest that SSX-C promotes PRC1.1 protein levels in part by stabilizing their presence on chromatin. Consistent with a SSX-C-mediated stabilization of PRC1.1, SSX-C expression strongly correlates with H2AK119ub1 levels in a manner that is not observed with eGFP or SSX-C mutants **(Fig. 6d, Extended Fig. 6a, d)**. Strikingly, by increasing PRC1.1 stability and H2AK119ub1 levels, SSX-C overexpression also impacts SS18 levels, which serve as a proxy for SS18-SSX1. This indicates that SSX-C alone is able to reinforce fusion binding **(Fig. 6e)**. Together, these results indicate that SS18-SSX promotes PRC1.1 activity via transcriptional and SSX-C-mediated mechanisms.

**Figure 6:**
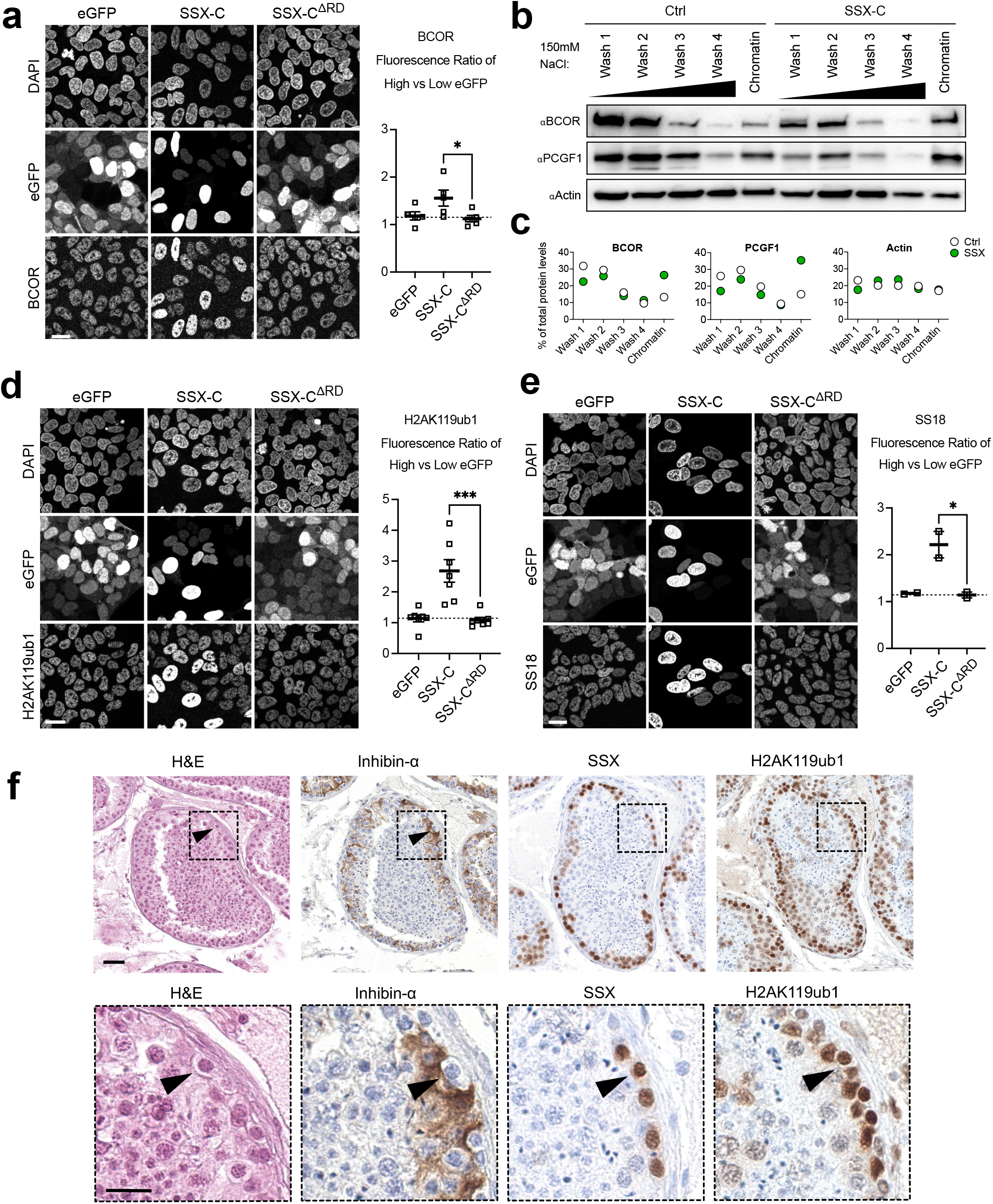
SSX-C increases PRC1.1 stability thus reinforcing H2AK119ub1 levels and SS18-SSX occupancy. **a)** Left, Immunofluorescence against BCOR in HS-SY-II synovial sarcoma cells expressing eGFP-fused constructs (eGFP, eGFP-SSX-C and eGFP-SSX-C^ΔRD^). Right, quantification of BCOR fluorescence ratio in high versus low eGFP cells in 5 biological replicates. Data represents the mean and the S.E.M. Asterisks represent p-values of paired one-tailed t-test between groups (* p< 0.05). Scale bars correspond to 20μm. **b)** Sequential chromatin washes assay using 150mM salt buffer in untransduced control (Ctrl) or eGFP-SSX-C expressing HEK293T cells. BCOR, PCGF1 or Beta-Actin as a loading control were detected by western blot. **c)** Quantification of the protein distribution for BCOR, PCGF1 or Beta-Actin in the various washes. Data represents the percentage of total protein levels. **d)** Left, Immunofluorescence against H2AK119ub1 in HS-SY-II cells expressing the indicated eGFP-fused constructs. Right, quantification of H2AK119ub1 fluorescence ratio in high versus low eGFP cells in 7 biological replicates. Data represents the mean and the S.E.M. Asterisks represent p-values of paired one-tailed t-test between groups (*** p< 0.001). Scale bars represents 20μm. **e)** Left, Immunofluorescence against SS18 in HS-SY-II cells expressing the indicated eGFP-fused constructs. Right, quantification of the SS18 fluorescence ratio in high versus low eGFP cells in 2 biological replicates. Data represents the mean and the standard error of the mean (S.E.M). Asterisks represent p-values of paired one-tailed t-test between groups (* p< 0.05). Scale bars correspond to 20μm. **f)** H&E and immunohistochemical staining for Inhibin-a, SSX and H2AK119ub1 in human testis. Scale bar correspond to 40μm in the upper panel. Lower panel is a close-up from the images shown in the upper panel (area marked by a dashed line) where the scale bar corresponds to 20μm.

To explore the role of wild-type SSX, we investigated whether SSX1 levels are associated with H2AK119ub1 in human testis where the protein is normally expressed. Publicly available single-cell RNA sequencing data from human testis shows that SSX1 is expressed in spermatogonial stem cells, differentiating spermatogonia as well as in early and late spermatocytes, but not in other testicular cell types **(Extended Fig. 6e)**^71^. Immunohistochemical staining of human testis revealed that H2AK119ub1 levels are not homogeneous, rather they are particularly high in cells around the outer edge of the seminiferous tubules, next to the basal lamina that correspond to spermatogonia (Inhibin-a negative cells) where SSX1 is also specifically detected **(Fig. 6f)**. These results suggest that the physiological role of SSX proteins is also linked to PRC1.1 function.

### A positive feedback loop promotes H2AK119ub1 levels during synovial sarcoma tumourigenesis

The previous *in vitro* results uncovered an autoregulatory feedback loop between SS18-SSX and PRC1.1, where increased H2AK119ub1 levels reinforce SS18-SSX binding. In light of these observations, we next assessed if the SS18-SSX oncoprotein promotes H2AK119ub1 *in vivo*. We took advantage of a synovial sarcoma mouse model in which SS18-SSX2 expression is conditionally-induced in *Hic1*-positive mesenchymal progenitors^72,73^ **(Fig. 7a)**. Similar to our observations in cell culture, SS18-SSX-positive tumour cells (marked by GFP) specifically exhibit high levels of H2AK119ub when compared to normal muscle **(Extended Fig. 7a-c)**. Moreover, higher levels of H2AK119ub1 where clearly detected at earlier time points following SS18-SSX induction, as early as 5 weeks post-induction and with a steady increase that was concomitant with the time course of tumour formation **(Figure 7b-d)**. These results indicate that the SS18-SSX/H2AK119ub1 feedback loop precedes transformation and underlies synovial sarcoma tumourigenesis *in vivo*.

**Figure 7:**
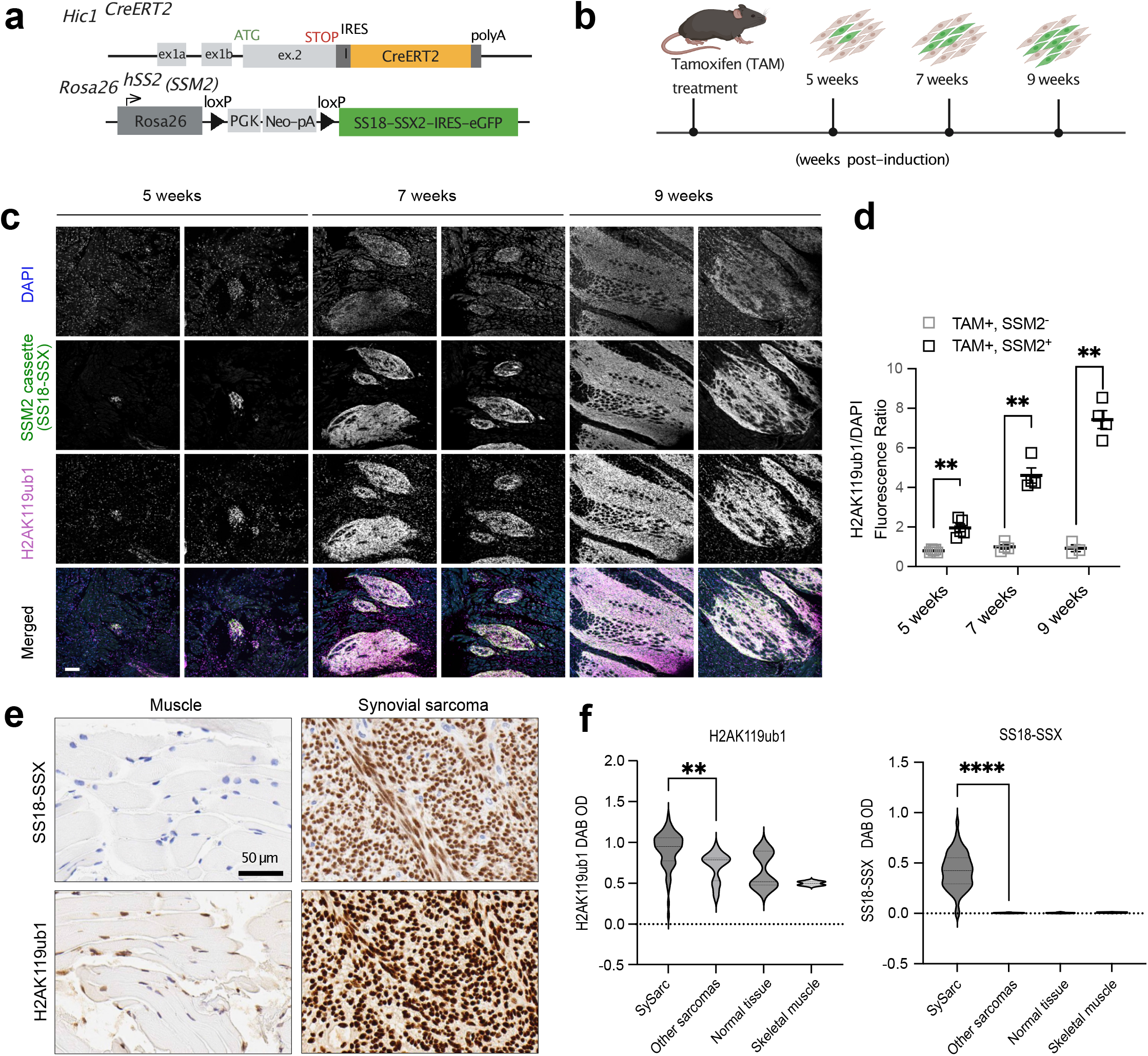
High levels of H2AK119ub1 are acquired during synovial sarcoma development. **a)** Overview of the *Hic1*^*CreERT2*^ knock-in allele (Scott et al, 2019) and of the *Rosa26*^*hSS2*^ (also known as SSM2) allele (Haidar et al, 2007) for conditional induction of SS18-SSX2 in Hic1-expressing mesenchymal progenitors. Upon tamoxifen treatment CreERT2 mediates recombination between the two LoxP sites in SSM2 mice, thereby removing the transcriptional stop signal and allowing transcription of SS18-SSX2-IRES-EGFP from the endogenous ROSA26 promoter. **b)** Illustration of the time line for the tissue sample collection of samples analysed in (c, d) 8-week-old mice were treated with tamoxifen and tongue muscle tissues were collected at 5, 7 and 9 weeks post-induction. Figure 7a and 7b were created with BioRender.com. **c)** Immunofluorescence of *Hic1*^*creERT2/creERT2*^; *Rosa26*^*SSM2/SSM2*^, mice tongue tissue at 5, 7 or 9 weeks after induction by tamoxifen treatment. The cells are stained for DAPI, SSM2 (eGFP) and H2AK119ub1. The scale bar represents 100 μm. **d)** Quantification of H2AK119ub1 signal intensity in normalised to DAPI signal intensity in 5 or 4 biological replicates of mice treated with tamoxifen (TAM) in SSM2 (GFP) negative tongue muscle or in adjacent SSM2 positive cell clusters. Asterisks represent p-values of paired one-tailed t-test between groups (** p< 0.01). **e)** limmunohistochemical staining for H2AK119ub1 on a tissue microarray of human surgical excised tissue specimens (left: skeletal muscle; right: synovial sarcoma). Scale bars correspond to 50μm in the left panel. **f)** Quantification of H2AK119ub DAB signal intensity across 37 synovial sarcomas (sample cores in duplicate), other sarcomas (1 case each of epithelioid sarcoma, sarcomatoid mesothelioma, Ewing sarcoma, sarcomatoid renal cell carcinoma, clear cell sarcoma, dedifferentiated liposarcoma, and myxoid liposarcoma) and normal tissues (normal skeletal muscle, ovarian stroma, breast glandular tissue, and testis controls). Quantification for the two skeletal muscle samples is also shown separately in the graph. All samples were stained in parallel on the same formalin-fixed, paraffin embedded tissue microarray slide. Asterisks represent p-values of Mann-Whitney test between groups (** p< 0.01).

We reasoned that if this autoregulatory feedback loop plays a role in human sarcomagenesis, increased levels of H2AK119ub1 would be a feature of human synovial sarcomas. To test this, we performed H2AK119ub immunohistochemistry on a synovial sarcoma tissue microarray of 37 patient samples. H2AK119ub1 exhibited stronger nuclear staining in synovial sarcomas in comparison to other sarcomas and normal tissues including skeletal muscle **(Fig. 7e, f)**. These results suggest that SS18-SSX activity in human synovial sarcoma also relies on an autoregulatory feedback loop that translates into high levels of H2AK119ub1 in these tumours. Taken together, our results uncover a feedback loop in which SS18-SSX/SSX-C and H2A ubiquitination cooperate to reinforce SS18-SSX presence on the chromatin and therefore its oncogenic function.

## DISCUSSION

Our study addresses the molecular mechanism underlying SS18-SSX chromatin recruitment and binding in synovial sarcoma. We demonstrate that the most critical domain of SS18-SSX1 for synovial sarcoma cell maintenance is at the SSX C-terminus where only 34 amino acids drive binding and maintenance of the oncofusion on chromatin. This highlights the critical role of the SSXRD domain in the precise recruitment of SS18-SSX at specific synovial sarcoma gene targets and indeed the SSX1 tail alone is able to reproduce SS18-SSX1’s genome wide occupancy in a SSXRD-dependent manner. This is consistent with the finding that some synovial sarcomas harbour translocations in which SSX1/2 is fused to other alternative partners such as EWSR1 and MN1^46^. Such occurrences suggest that the SSXRD domain, by mediating recruitment of transcriptional activators to induce Polycomb target genes during sarcomagenesis, is the key determinant of a synovial sarcoma signature and that direct de-regulation of the mSWI/SNF complex through SS18 is not essential to all synovial sarcomas.

We also show that SSXRD binds chromatin environments rich in H2AK119ub1 and histones MacroH2A. Intriguingly, SSXRD is also present in PRDM9, a DNA binding protein critical in delineating homologous recombination hotspots during meiosis via H3K4me3 deposition^74–76^. Recent work showed that removing SSXRD in PRDM9 led to a complete loss of H3K4me3 deposition at PRDM9 hotspots^77^ potentially suggesting a role of SSXRD in driving PRDM9 chromatin localisation as well.

Our data supports the idea that H2AK119ub1 is important for SS18-SSX specific chromatin targeting^48^, and further shows that in synovial sarcoma, PRC1.1 is central in establishing H2AK119ub deposition and therefore oncofusion protein occupancy and maintenance. Hence, PCGF1 removal leads to a global erosion in SS18-SSX binding. These results support a role for PRC1.1 as the main contributor of genome-wide H2AK119ub1 deposition as observed in mouse embryonic stem cells^10^, and suggest that other variant PRC1 complexes may have alternative roles in synovial sarcoma. Moreover, we show for the first time that PRC1.1 activity can deposit histones MacroH2A alongside H2AK119ub1. Removing MacroH2A did not impact H2AK119ub1 deposition indicating that deposition of the histone variant occurs downstream of the establishment of the histone mark. Interestingly, in synovial sarcoma cells, this occurs independently of PRC2 which was so far the only complex thought to regulate MacroH2A deposition^65^.

Our data also revealed a strong interplay between SS18-SSX and PRC1.1 activity leading to a positive feedback loop that results in increased H2AK119ub1 in murine and human synovial sarcomas. Two distinct mechanisms mediate this interplay. On one hand SS18-SSX binds to and positively regulates the transcriptional level of PRC1.1 genes, *BCOR* and *RYBP*. On the other hand, the SSX C-terminus induces an increase in H2AK119ub1 by stabilizing the PRC1.1 complex levels and chromatin binding. Both simultaneously act to further promote SS18-SSX presence on the chromatin **(Fig. 8)** thereby reinforcing its oncogenic activity. This model is in agreement with a previous study showing that RYBP occupancy is induced by SS18-SSX expression in murine mesenchymal stem cells^78^. The feedback loop we identified is also reminiscent of the role of RYBP in the PRC1 complex, where it both promotes interactions within the complex leading to increased complex stability^70^ and recognizes and binds H2AK119ub1-modified nucleosomes to further promote H2AK119ub1 deposition^79^. This work further sheds light onto the central role that PRC1 activity, and is derivate H2AK119ub1 histone mark, plays in driving and sustaining synovial sarcoma and supports inhibition of PRC1.1 as a potential therapeutic strategy.

**Figure 8:**
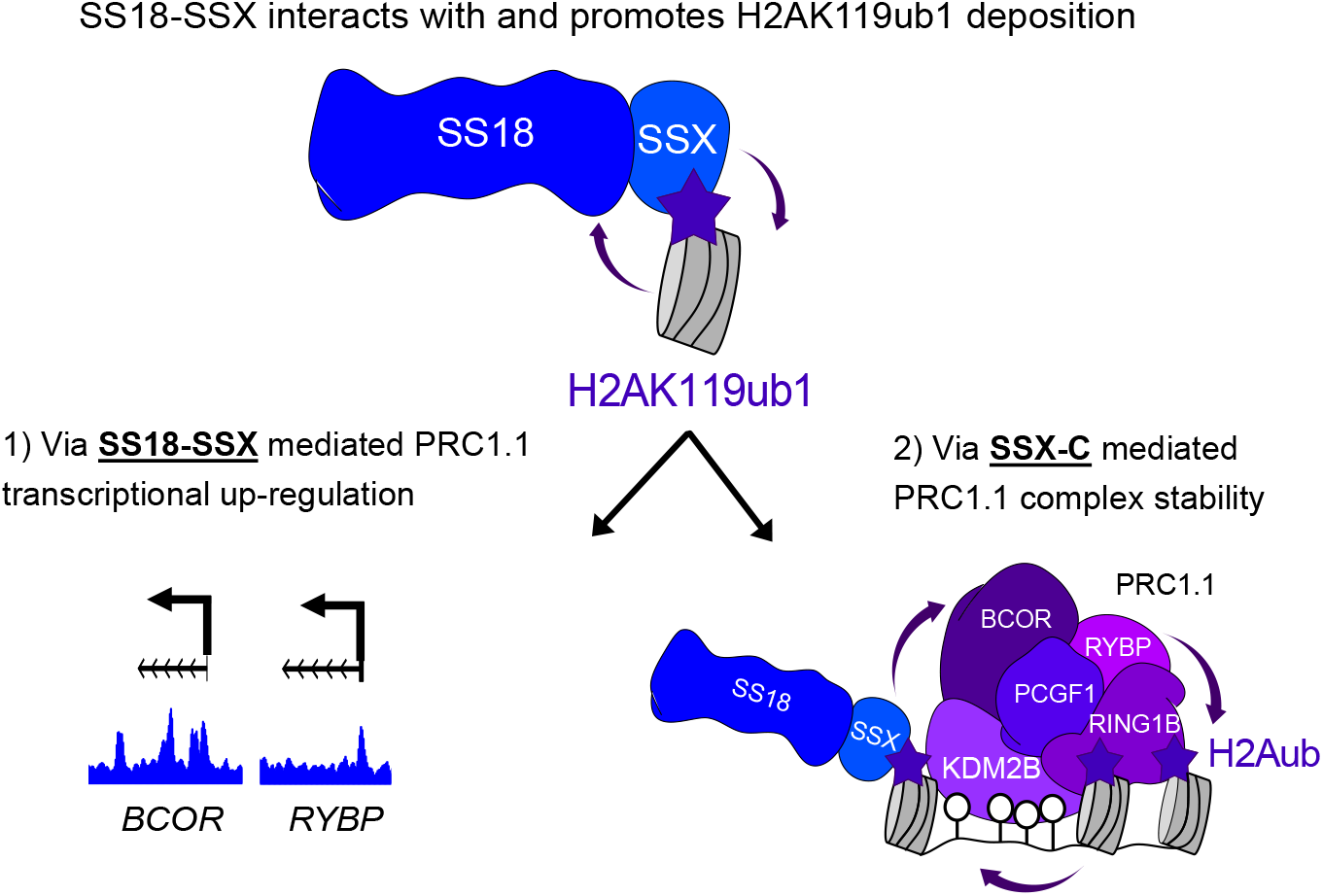
An autoregulatory feedback loop converging on H2AK119ub1 drives synovial sarcoma. Model depicting the strong interplay between SS18-SSX and H2AK119ub1 where SS18-SSX interacts with histones H2A which are ubiquitinated on their lysine K119. SS18-SSX then promotes further levels of H2AK119ub1 via two distinct mechanisms: 1) by stimulating the transcription of PRC1.1 members *BCOR* and *RYBP* as direct targets of the fusion and 2) by increasing the stability of the PRC1.1 complex on chromatin. In both cases H2AK119ub1 levels increase and therefore reinforce SS18-SSX’s presence on chromatin.

These findings are also important in light of a putative role of full length wild-type SSX family proteins, which have been reported to be expressed in synovial sarcomas^32,80^, in further promoting oncofusion protein activity. Moreover, SSX proteins are cancer-testis antigens that are abnormally present in various cancers such as melanoma, breast and prostate cancer^81,82^. Therefore, the interplay between SSX and H2AK119ub1 could impact a wider range of other malignancies.

Our study describes a central role for H2AK119ub1 in driving synovial sarcoma and in doing so it highlights a key role for PRC1 beyond cell fate decisions and development, which is further supported by the occurrence of main driving genetic events involving *BCOR* in several pediatric tumours^24–29^. Further studies will uncover to what extent “PRC1-dependent” tumours share molecular characteristics and circuitries that can be exploited therapeutically.

## METHODS

### Cell Culture

Human synovial sarcoma cell lines: HS-SY-II (RRID:CVCL_8719)^83^ and SYO-1 (RRID:CVCL_7146)^84^ were obtained fom their original source laboratories. Human osteosarcoma KHOS-240S (RRID:CVCL_2544) and Human Embryonic Kidney HEK293T (RRID:CVCL_0063) were purchased from the American Type Culture Collection (ATCC). Cells were cultured in DMEM (Gibco) supplemented with 10% Fetal Bovine Serum. ASC52telo, hTERT immortalized adipose derived Mesenchymal stem cells were purchased from ATCC (SCRC-4000) and were cultured in MesenPRO RS™ Medium (Gibco, 12746-012) supplemented with L-glutamine (Sigma-Aldrich, G7513-100ML) at a final concentration of 2 mM.

### Virus Production and transduction

For lentivirus production, 1×10^6^ HEK293T cells were transfected with 3μg constructs and helper vectors (2.5μg psPAX2 and 0.9μg VSV-G). For retroviral infection, 10×10^6^ 293T-gag-pol cells were transduced with 20 ug of MSCV vectors and 2.5ug of VSV-G. Transfection of packaging cells was performed using polyethyleneimine (Polysciences, 23966-2) by mixing with DNA in a 3:1 ratio. Viral supernatants were collected 48h after transfection, filtered through a 0.45 μm filter and supplemented with 4 μg /ml of polybrene (Sigma) before adding to target cells. Downstream experiments using sgRNAs knockout were performed 10 days after sgRNA induction (CUT&RUN, RNA sequencing) or 12 days after knockout (Immunofluorecence). Downstream experiments using eGFP or MBD constructs (Salt Extraction, Imaging, Nuclear co-IP) were performed 6 days after induction.

### Generation of Cas9 stable cell lines

For stable expression, HS-SY-II and SYO1 synovial sarcoma cell lines were transduced with lentiCas9-Blast^85^ (Addgene, #52962) and selected using 20 μg/ml of blasticidin to generate stable Cas9-expressing cell lines. Cells were consequently transduced with sgRNAs and selected with puromycin. For inducible expression HS-SY-II and SYO1 were transduced with a lentivirus expressing the reverse tetracycline-controlled transactivator (rtTA3), the ecotropic receptor (EcoRec) and a hygromycin resistance gene (pRRL-SFFV-rtTA3-IRES-EcoR-PGK-hygro, a gift from Johannes Zuber). Transduced cells were selected with 40 μg/ml hygromycin (Invitrogen). Selected cells were next transduced with a lentivirus expressing Cas9 and BFP (Blue Fluorescent Protein) under regulation of tetracycline-responsive element promoter (pRRL-TRE3G-Cas9-P2A-BFP, a gift from Johannes Zuber). Cas9 and BFP expression induction was achieved by treatment of the cells for 3 days with doxycycline (Dox) (1µg/ml) (Hyclate, Alfa Aesar) and single cell clones were seeded by sorting BFP positive cells using the BD FACSAria™ Cell Sorter system. To measure Cas9 efficient and leakiness, HS-SY-II and SYO1 Dox inducible Cas9 single cell clones were transduced with a sgRNA against the surface molecule CD46 (pLKO-U6sgRNA_CD46_improved-EF1s-GFP-P2A-Puromycin) (sgRNA CD46: 5’-GGAGTACAGCAGCAACACCA-3’) as described in^86^. Inducible Cas9 expression was activated for 6 days using doxycycline to remove SSX or PCGF1 when applicable.

### Plasmid Cloning

sgRNA for CRISPR knock-out were designed using Sanjana lab tool (http://guides.sanjanalab.org/) and cloned as previously described^38,85^ (see Table S1 for sgRNA sequences). Briefly, sgRNAs were cloned by annealing two DNA oligos and ligating into a BsmB1-digested pLKO1-puro-U6-sgRNA-eGFP. Transformation was carried into Stbl3 bacteria. eGFP constructs were cloned into pLV-EF1a-IRES-Neo lentiviral backbone^87^ (Addgene, #85139) containing a Neomycin selection cassette. cDNAs were adapted from the MSCV-HA-PGK-Puro plasmids ^23^. For the SSX fusion vectors, cDNA of EWSR1 and MN1 were obtained from the DKFZ cDNA clone repository and assembled with an HA-tag at the N-terminus and SSX at the C-terminus into a MSCV-PGK-Puro backbone in a single step using NEBuilder HiFi DNA Assembly (NEB, #E2621).

NanoBRET plasmids pHTN-HaloTag-CMV-neo (Promega, G7721) and pNLF1-N-CMV-Hygro (Promega, N1351) were obtained from Promega. Histone H2A cDNA were PCR amplified from pCDNA3.1-Flag-H2A and pCDNA3.1-Flag-H2A K118-119R^51^ (Addgene, #63560, #63564). Histones Macro-H2A cDNAs were obtained for the DKFZ cDNA clone repository.

MBD-V5 constructs were cloned into pLV-EF1a-IRES-Neo. Luciferase was amplified by PCR from pT3-EF1a-NrasG12V-GFP-P2A-Luc2 (a gift from Scott Lowe’s laboratory), KDM2B was amplified from pUC19-hKDM2B (Sino Biological, HG20918-U) and the ZF-CxxC mutant was generated with PCR using mismatched primers (Q5). The MBD sequence was amplified using pENTR-MBD1^88^ (Addgene, #47057) as template. The assembly was designed and performed in a single step adding the MBD, the cDNAs and V5-NLS using NEBuilder HiFi DNA Assembly.

### Cell Competition Assays

HS-SY-II and KHOS-240S Cas9 cells were transduced with an empty plasmid (Empty Vector), plasmid containing sgRNA targeting PCGF1, or sgRNA targeting both MacroH2A. Infections were done with a virus dilution of 1:10 to obtain an infection efficiency of around 70-80%. Infected cells become GFP+ due to the backbone of the sgRNA. The cells were then cultured over a period of 25 days, and the percentage of GFP+ cells measured using a Fortessa FACS machine. Data was analysed using Flowjo software.

### CRISPR/Cas9 gene-tilling screen

sgRNAs library cloning and screen deconvolution were performed as previously described^89,90^. Briefly, sgRNAs targeting the entire coding sequence of SS18 and SSX1 were designed using http://crispr.mit.edu/ and Benchling (https://benchling.com) and cloned into pLKO-U6-sgRNA-improved-EF1s-GFP-P2A (gifted by Darjus F Tschaharganeh). 211 sgRNA were designed spanning the length of the isoform 1 of SS18 (NT 010966) and 90 sgRNA targeting isoform 1 of SSX1 (NT 011568). Additionally, 200 safe sgRNA were added as negative controls; these guides target the non-genic region of genome^91^. Stable Cas9-expressing cell lines were transduced to about 30% efficiency. After 3 days of infection, cells were selected with 2µg/ml of puromycin. Cells were passaged with the number of cells kept at 3000 times the size of the library, i.e., at least 1.56×10^6^ cells were passaged. After 15 population doublings the cells were harvested, and their genomic DNA was extracted using phenol extraction method. The region spanning the sgRNA was amplified via using custom primers. Amplicons were sent for next generation sequencing using NextSeq 550 SR 75 HO. Files were demultiplexed and counts were mapped on the library using Mageck tool. To identify individual regions which are more important for cell survival, ProTiler tool was used to identify CKHS (CRISPR Knockout Hyper-Sensitive) regions were identified.

### Live Imaging

30 000 HEK293T cells transduced with the various eGFP constructs were seeded in an 8 well chamber slide (µ-Slide 8 Well high, Ibidi). Cells were then imaged within the next 48h using Leica TCS SP5 inverted Confocal with the HCX PL APO 63x/1.40-0.60 Oil Lbd BL objective, a single z-stack was captured. DNA was stained 30 minutes prior image acquisition using NucBlue Live ReadyProbes Reagent (Hoechst 33342) (Invitrogen).

### Immunofluorescence staining

Between 0.5×10^6^ to 1×10^6^ cells were seeded 6 days after induction in 6-well plates containing coverslips. Cells were fixed the following day with 4% paraformaldehyde for 10 minutes. Permeabilization was performed using TritonX (0.1 % in PBS) for 12 min followed by incubation with blocking solution (1% BSA, 0.1% Gelatin Fish in PBS) for 1 hour. Incubation with the primary antibody was performed in blocking buffer for 1 hours at RT. Cells were washed, incubated with secondary antibodies for 1h and mounted in Vectashield Antifade Mounting Medium containing DAPI (Vectorlabs). For 4-color immunofluorescence using V5-555 antibody (Invitrogen), after the secondary antibody, cells were washed and incubated with V5-555 for 1h prior to mounting. Antibodies used are listed in Table S2.

### Image Capture and Analysis

Confocal images were acquired on Leica TCS SP5 inverted Confocal using the HCX PL APO 63x/1.40-0.60 Oil Lbd BL objective, a single z-stack was captured. Samples were imaged using 405, 488, 561, 594 and 633nm laser lines using sequential mode in the Leica Application Suite software. For illustration, samples were imaged using a 512×512 format at a 100Hz speed using line averaging at 4 with a zoom factor of 11. Images were then smoothed and adjusted for brightness and contrast using the ImageJ/Fiji software. For MBD foci quantifications, images were acquired using a 512×512 format at a 700Hz speed with a zoom factor of 1.7. Between 50 and 100 foci were counted per replicate, each MBD foci was selected, and only co-occurring foci were counted.

For H2AK119ub, BCOR and SS18 signal intensity quantifications in eGFP induced cells, nuclei were detected and selected using Li’s thresholding on ImageJ/Fiji software. Signal intensity for each selected nuclei was measured for all the channels (405-DAPI, 488-eGFP, 594-H2AK119ub /BCOR/SS18 and 647-H2AK119ub when applicable). The corrected mean intensity of the 488, 594 and 647 channel was calculated by dividing the mean signal intensity of each nucleus by its corresponding mean DAPI intensity to normalize the signals. eGFP signals were separated in two groups, low eGFP (corrected mean intensity <1) and high eGFP (corrected mean intensity > 1). In the low and high eGFP group, the average of the corrected mean intensity of the 594 and 647 channels was calculated. Finally, the ratio of the high versus low was used to display the change in signal intensity in the high eGFP population (Average of the corrected mean intensity _high eGFP /_ Average of the corrected mean intensity _low eGFP_). For each biological replicates, between 50 and 250 nuclei were analyzed.

### Mouse Model for conditional SS18-SSX2 expression

The mouse model of synovial sarcoma used herein (Scott et al., unpublished data) is based on the hSS2 model with a conditional SS18-IRES-eGFP allele knocked into the Rosa26 locus^72,73^. Animals were maintained and experimental protocols were conducted in accordance with approved and ethical treatment standards of the Animal Care Committee at the University of British Columbia.

#### Tissue processing and staining

To enable detection of native eGFP expression in processed tissue samples, mice at clinical endpoint were humanely euthanized by intraperitoneal injection of Avertin (400 mg/kg) and the tongues were removed. Wild type tongue samples were obtained from age matched Cre negative control animals. Dissected tongues were immersed in 2% paraformaldehyde fixative for 48 hrs at 4°C. Samples were then washed 3 × 30 mins in PBS and then immersed through a gradient of sucrose solutions from 10%-50% at 4°C for > 4 hrs each before being embedded in cryomolds (Polysciences 18646A) using OCT (Sakura Finetek, 4583) and frozen in an isopentane bath cooled by liquid nitrogen. Cryosections were cut (Leica CM3050S) at a thickness of 20 um and mounted onto Superfrost Plus slides (VWR 48311-703). Slides were thawed at 37°C for 30 mins, washed 3 × 10 mins in PBS and incubated for 1 hr in PBS containing 10 mg/mL sodium borohydride (Sigma 213462) to quench autofluorescence. Following this treatment, slides were briefly washed with PBS and incubated in block solution containing 2.5% BSA (Sigma A7030) and 2.5% Goat serum (Gemini 100-190) for 90 min at room temperature prior to incubation in primary antibody dissolved in block solution (1:100), overnight at 4°C. Primary antibody solution was removed and slides were washed 3 × 5 mins in PBS before Alexa Fluor conjugated secondary antibodies were applied to the slides for 45 min. After secondary antibody incubation, 3 × 5 min PBS washes were performed and sections were counterstained with DAPI (600 nM in PBS) for 5 mins, rinsed and mounted with Aqua Polymount (Polysciences 18606).

#### Image acquisition and quantification

Confocal images were collected using a Nikon Ti-E inverted microscope with an A1R HD25 confocal scanning head and acquired in Nikon Elements software. For quantification, a single z-stack was selected and the image was first smoothed. Nuclei were detected using Li’s thresholding on ImageJ/Fiji software. Signal intensity for each selected nuclei was measured for the channels 405-DAPI, 488-eGFP (SSM2 cassette) and 647-H2AK119ub. The ratio intensity of H2AK119ub1 over DAPI was calculated by dividing the 647 mean signal intensity over its corresponding 405 mean signal intensity.

### Human testis immunohistochemical imaging

For immunohistochemical analyses, formalin-fixed and paraffin embedded (FFPE) tissue samples of non-neoplastic human testis were retrieved from the archives of the Institute of Pathology, University Hospital Heidelberg, Heidelberg, Germany. Use of patient samples was approved by the ethics committee of the University of Heidelberg (S-442/2020). Four μm sections were cut and mounted on SuperFrost Plus Adhesive slides (Thermo Scientific), followed by deparaffinization and heat induced antigen retrieval (97 degree celcius) in high pH buffer (pH 9) for 30 minutes. Primary monoclonal mouse antibodies for Inhibin α (ready to use, clone R1, Dako Omnis, Agilent, Glostrup, Denmark), SSX (dilution 1:100) and H2A119ub1 (dilution 1:500) listed in Table S2 were each incubated for 25 minutes. Visualization was performed using the ready to use Polyview Plus HRP (anti-mouse) reagent (Enzo Life Sciences, Farmingdale, USA). Sections were counterstained with hematoxylin.

### Human synovial sarcoma tissue microarray immunohistochemical imaging

H2AK119Ub and SS18-SSX immunohistochemistry were performed on a 4 µm section of a formalin-fixed, paraffin-embedded human tissue microarray (TMA) consisting of: 37 synovial sarcoma cases; 1 case each of epithelioid sarcoma, sarcomatoid mesothelioma, Ewing sarcoma, sarcomatoid renal cell carcinoma, clear cell sarcoma, dedifferentiated liposarcoma, and myxoid liposarcoma, as well as normal skeletal muscle, ovarian stroma, breast glandular tissue, and testis controls from Vancouver General Hospital. Cases were included as 0.6 mm patient sample cores in duplicate. The assays were run with the following conditions, via a Leica Bond RX (Leica Biosystems, Buffalo Grove, IL, USA). Heat-induced epitope retrieval was performed using citrate-based BOND Epitope Retrieval Solution 1 (Leica Biosystems, Buffalo Grove, IL, USA) for 10 min, 10 min, and 20 min, respectively. The primary antibodies H2AK119Ub (Cell Signaling Technology, #8240, Danvers, MA, USA), and SSX-SS18 (Cell Signaling Technology, #72364S, Danvers, MA, USA) were incubated at 1:400 for 30 min and 1:300 for 15 min, respectively, at ambient temperature. Staining was visualized using the BOND Polymer Refine Detection kit (Leica Biosystems, DS9800, Buffalo Grove, IL, USA), which includes a 3,3’-diaminobenzidine (DAB) chromogen and hematoxylin counterstain. TMA virtual slide scans were then generated on a Leica Aperio AT2 (Leica Biosystems, Buffalo Grove, IL, USA) at 40x magnification. Each individual patient sample core was analysed using HALO and HALO AI (Indica Labs), which required user annotated training data to develop an artificial intelligence segmentation network for nuclear identification. The TMA module was implemented to extract individual patient core images from the TMA whole slide scan. The Multiplex IHC module was trained to identify DAB staining using representative pixels for delineation from hematoxylin in order to determine average DAB nuclear optical density.

### NanoBRET

NanoBRET Protein:Protein Interaction assay was performed following the manufacturer’s conditions (Promega, N1662). 0.5×10^6^ HEK293T cells were plated the day before transfection in a 12 well plate. 2µg of HaloTag plasmid (empty, SS18, SS18-SSX, SSX-C, SSX-C^ΔRD^ or SSX-C^E184*^) + 0.2µg of NanoLuc plasmid (H2A WT, H2A^K118K119R^, MacroH2A1.2 or MacroH2A2) were transfected using polyethylenimine by mixing with DNA in a 3:1 ratio. 48h after transfection, cells were counted and adjusted to a final concentration 2×10^6^ cells/ml. Cells were passed in a 96-well white plate. For each condition, 10µl (20000 cells) were seeded in 4 different wells. Each well was supplemented with 90µl of Opti-MEM I Reduced Serum Medium, no phenol red (Gibco, 11058-021) containing 4% FBS with either 100nM HaloTag NanoBRET 618 Ligand (+ ligand, experimental samples in 2 technical replicates) or 0.1% DMSO final concentration (– ligand, no-acceptor controls in 2 technical replicates). The next day, 72h post transfection, 25µl of 5X NanoBRET Nano-Glo in Opti-MEM I Reduced Serum Medium was added on all the wells. Measurements of NanoBRET bioluminescent donor emission (460nm) and acceptor emission (618nm) were performed within 10 minutes of substrate addition using a PHERAstar Microplate Reader (BMG LABTECH) using 450nm and 620nm filters. NanoBRET calculations were done using the followings steps: the raw NanoBRET ratio (BU) was obtained by dividing the acceptor emission value (620nm) by the donor emission value (450nm) for each sample. BU were then converted to milliBRET units (mBU) by multiplying each raw BRET value by 1000. The final BRET ratio (mBU) displayed in the figures is calculated for each biological replicate by subtracting the mean of the two experimental replicates (+ ligand) with the mean of the two no-ligand control replicates (-ligand).

### Chromatin Salt Extraction/Sequential chromatin washes

Chromatin salt extraction was adapted from Herrmann et al^44^. Approximatively 10×10^6^ cells were harvested and washed twice in PBS. Cell pellets were then washed in a series of chromatin salt extraction buffers containing 0.1% Triton X, 300mM Sucrose, 1mM MgCl2, 1mM EGTA, 10mM PIPES and NaCl at increasing concentrations: 80mM, 150mM, 300mM or 500mM. All buffers were supplemented with protease inhibitors (Protease inhibitor tablets, Roche). Cell pellets were resuspended and incubated in 50μl of chromatin salt extraction buffer for 10 min at room temperature and pelleted at 2000g for 5 min. The supernatant was transferred to a new tube and supplemented with 2X Laemmli (Invitrogen) and kept on ice after a 5-minute denaturation step at 95°C. For the chromatin extraction after the last 500mM wash, pellets were resuspended in 500mM NaCl chromatin salt extraction buffer supplemented with 2X Laemmli. The chromatin sample was then denatured at 95°C for 5 minutes and sonicated. Chromatin samples were then centrifuged at full speed for 5 minutes to get rid of the DNA debris and transferred to a new tube. Sequential chromatin washes were performed similarly but the cells and the chromatin were washed at constant salt concentration of 150mM for 4 washes, the chromatin fraction was then sonicated as above. Samples were then used for Western Blotting. The signal intensity in the various salt fraction was measured using the maximum intensity of a square containing the band in the ImageJ/Fiji software. The total protein level was calculated using the sum of the maximum intensity as a proxy. Each intensity/salt fraction was then represented as a percentage of total protein levels.

### Histone Acid Extraction

Approximatively 1×10^6^ cells were harvested and washed twice in PBS. Cells were resuspended in 100µl PBS + 0.5% Triton-X and incubated for 10 min on ice. After centrifugation at 6,500g for 10 min at 4°C, nuclei were washed a second time in 100µl PBS + 0.5% Triton-X. Nuclear pellets were then resuspended in 25µl of 0.2 N HCl. Histones were released overnight at 4°C and DNA debris pelleted at 6,500 g for 10 min at 4°C. Histone acid extracts were neutralised with 2.5µl of 2M NaOH. After 2X Laemmli addition and denaturation at 95°C for 5 minutes, samples were loaded onto a Western Blot gel.

### Whole cell protein extracts & Western Blotting

Cells grown in 6-well plates were harvested and washed in PBS. Cell pellets were incubated with RIPA buffer (Cell Signaling) supplemented with protease inhibitors (Roche) for 30 min and cleared by centrifugation (15 min 14.000 rpms 4C). Protein lysates were quantified using a BCA protein assay (Pierce). Lysates were then denatured in 2X Laemmli at 95°C for 5 minutes and run in Mini-PROTEAN Precast Gels (BioRad) and transferred onto membranes using Trans-Blot Turbo. Membranes then were blocked in 5% milk in TBST. Western were visualized using Amersham Imager 680.

### Nuclear Immunoprecipitation

Approximatively 5×10^7^ cells were harvested and washed twice in PBS. Nuclei isolation, nuclear fraction digestion and collection was performed using Nuclear Complex Co-IP Kit (Active Motif, 54001). 25µl per IP of GFP-Trap Magnetic Agarose beads (Chromotek) were washed twice in 1X IP Low buffer supplemented with protease inhibitor and PMSF following manufacturer’s guidelines (Active Motif, 37511). 200µl of nuclear extracts were incubated with the GFP-Trap beads for 1hour at 4°C. Beads were then washed three times in 1X IP Low buffer and resuspended in 100µl of 1% SDS. Beads were denatured at 95°C for 5 minutes and the supernatant was submitted for mass spectrometry at the EMBL Proteomics Core Facility. Data analysis was performed by the Facility. The raw output files of IsobarQuant (protein.txt – files) were processed using the R programming language. Only proteins that were quantified with at least two unique peptides were considered for the analysis. Raw signal-sums (signal_sum columns) were first cleaned for batch effects using limma^92^ and further normalized using variance stabilization normalization^93^. Different normalization coefficients were estimated for control conditions in order to maintain the lower observed abundance.

### Chromatin Immunoprecipitation (ChIP)

HA-tag ChIP on HS-SY-II cells expressing MSCV-HA-eGFP-PGK-Puro, MSCV-HA-eGFP-SSX-C-PGK-Puro or MSCV-HA-eGFP-SSX-C^ΔRD^-PGK-Puro was performed following puromycin selection and collected 6 days following transduction. HS-SY-II cells were pre-fixed for 20 minutes with 1.5mM ethylene glycol bis(succinimidyl succinate) (Thermo Scientific) and then fixed with 1% formaldehyde for 15min and the cross-linking reaction was stopped by adding 125mM of glycine. Cells were washed twice with cold PBS and lysed in swelling buffer (150mM NaCl, 1%v/v Nonidet P-40, 0.5% w/v deoxycholate, 0.1% w/v SDS, 50mM Tris pH8, 5mM EDTA) supplemented with protease inhibitors. Cell lysates were sonicated using a Covaris E220 Sonicator to generate fragments less than 400 bp. Sonicated lysates were centrifuged and incubated overnight at 4°C with HA-tag Abcam 9110. Immunocomplexes were recovered by incubation with 30ul protein A/G magnetic beads (Thermofisher) for 2h at 4°C. Beads were sequentially washed twice with RIPA buffer and finally TE buffer.

### Cleavage Under Targets and Release Using Nuclease (CUT&RUN)

SS18-SSX, H2AK119ub1 and MacroH2A2 occupancy was assayed using CUTANA ChIC/CUT&RUN Kit (EpiCypher, 14-1048) following manufacturer’s protocol. Briefly, 0.5 million human synovial sarcomas cells (HS-SY-II or SYO-I) were harvested per sample and bound to activated Concanavalin A magnetic beads. Beads were then incubated at 4°C overnight with 1:50 dilution of antibodies per sample. Chromatin digestion was performed for 2 hours at 4°C. Digestion is then stopped by chelating Ca++ ions in a buffer containing *E*.*coli* DNA for Spike-in. DNA Fragments are then released in solution after a 10 min incubation at 37°C on a ThermoMixer at 500 rpm. DNA Fragments were then purified using CUTANA DNA Purification Kit (EpiCypher).

### Library preparation

DNA fragments obtained after ChIP or CUT&RUN were quantified using Qubit dsDNA HS Assay Kit (Invitrogen). 5ng of DNA was used for library preparation using NEBNext Ultra II DNA Library Prep Kit for Illumina (NEB, E7645S), SPRIselect beads (Beckman Coulter, #B23317) and NEBNext Multiplex Oligos for Illumina (NEB, Set 1 #E7335S, Set 2 #E7500S). ChIP libraries were done following NEB’s guidelines (NEB, E7645S), CUT&RUN libraries were done following CUTANA ChIC/CUT&RUN Kit (EpiCypher, 14-1048) adapted protocol. ChIP libraries were sequenced as 75 bp Single-Read on Illumina NextSeq 550 platform High-Output. CUT&RUN libraries were sequenced as 75 bp Paired-End reads on Illumina NextSeq 550 platform Mid-Output.

### ChIP-seq Analysis

SS18-SSX1 (HA) input (SRR6451607), SS18-SSX1 (HA) IP (SRR6451595), KDM2B input (SRR6451587) and KDM2B IP (SRR6451586) were obtained from deposited GEO under the accession number GSE108926. Raw reads were trimmed for quality and Illumina adapter sequences using trim-galore, then aligned to the human genome assembly hg38 using Bowtie 2^94,95^ (with “--very-sensitive” option). ChIP signals were normalised to their respective inputs using the pileup function from MACS2^96,97^ using corresponding input for background normalization. To visualize ChIP-Seq tracks, normalized bigWig files were generated with ucsc-wigtobigwig tool. HA-SS18-SSX1 peaks (n=26805) were generated with the MACS2 function (with “--no model”, “--qvalue 0.05”, “--broad” options) and normalized to input.

### CUT&RUN Analysis

Paired-end reads were aligned to the hg38 and *E*.*coli* K12, MG1655 reference genome using Bowtie 2 (with options for hg38: --local --very-sensitive-local --no-unal --no-mixed --no-discordant --phred33 -I 10 -X 700 and for K12 --end-to-end --very-sensitive --no-overlap --no-dovetail --no-mixed --no-discordant --phred33 -I 10 -X 700). To internally calibrate our CUT&RUN experiments, we used the exogenous *E*.*coli* genome to quantitatively compare the genomic profiles as previously described ^98^. We first calculated the percentage of spike-in reads that align uniquely. We then normalized the sequencing depth (x% of spike-in reads in total reads) using the scaling factor so that *E. coli* spike-in signal is set to be equal to 1 across all samples: scaling factor = 1/ x% spike-in reads in total reads.

Genome coverage files were generated using bamCoverage^99^ with 50bp bins, no normalisation and scaled (--scaleFactor). When applicable, correlation of replicates was confirmed using deeptools functions multiBigwigSummary and plotCorrelation^99^. For peak calling, the MACS2 callpeak function was used on the aligned BAM files (with “--nomodel”, “--pvalue 0.001”, “--broad” options, “--keep-dup all”).

Heatmap of Spearman correlation coefficients from coverages were computed over CpG islands UCSC track from bigWig files using deeptools multiBigwigSummary and plotCorrelation^99^.

### ChIP and CUT&RUN Data Visualisation

Genome tracks were visualized using WashU Epigenome Browser. For heatmaps and metaplot profiles read densities of the various IPs were centered around SSX peaks (ChIP), HA-SS18-SSX1 peaks for HS-SY-II (replicates pooled together, n=52027) or SS18-SSX2 peaks for SYO-I (n=61940). A +/- 5 kb window from peak center was aligned and binned with 50 bp using computeMatrix and plotProfile/plotHeatmap functions from deeptools^99^.

### Human single cells testis Atlas

tSNE plots were obtained from the Human Testis Atlas Browser by Cairns Lab^100^. https://humantestisatlas.shinyapps.io/. Data was acquired on young adults 17, 24 and 25 years old.

### Data availability

HA-SS18-SSX1 and KDM2B ChIP sequencing data re-analysed in Figure 1 originates from GEO accession number GSE108929. The GEO accession number for the ChIP-Seq and CUT&RUN-Seq data reported in this paper is GSE205955.

## AUTHORS’ CONTRIBUITIONS

N.S.B. conceived the study, designed, performed, and analysed the experiments, and wrote the manuscript. V.D. generated the Cas9 cell lines, conducted the CRISPR/Cas9 screen, the competition assays and assisted with the CUT&RUN experiments. R.S.W and T.M.U provided the mouse model data. F.K.F.K. performed immunohistochemistry analysis of human testis. A.S, A.P, L.G., L.W. assisted in experiments and reagents production. F.J.S-R. assisted with CRISPR/Cas9 library cloning and screen deconvolution. M.T., S.T. and T.O.N. provided the analysis for the synovial sarcoma tissue microarrays. A.B. conceived and coordinated the study and wrote the manuscript. All authors read and approved the final manuscript for publication.

## ACKNOWLEDGEMENTS

We thank Wei He from Han Xu’s laboratory for help in running ProTiler analysis for the *SS18-SSX1* gene-tilling screen. Scott W. Lowe, Darjus Tschaharganeh, Johannes Zuber and Steven Henikoff for sharing reagents and protocols. Deepti Talwar from the Tobias Dick group at the DKFZ for assistance in measurements for the NanoBret assays. The EMBL Mass Spectrometry Facility members Mandy Rettel and Frank Stein for sample processing and data analysis. We also thank Robert Illingworth and Christopher Playfoot for their useful comments and discussion regarding the manuscript and members of the paediatric soft-tissue sarcoma lab and the U54 Synovial Sarcoma consortium for feedback and fruitful discussions. This project has received funding from the European Research Council (ERC) under the European Union’s Horizon 2020 research and innovation programme (grant agreement n° 805338) (A.B.) and from the National Institutes of Health/National Cancer Institute (NIH/NCI) U54CA231652 (T.O.N., T.M.U. and A.B.). T.O.N. and T.M.U. were additionally supported by grants from the Canadian Cancer Society (705615) and the Terry Fox Research Institute (1082). N.S.B. was supported by a DKFZ Postdoctoral Fellowship. F.J.S-R. was supported by the MSKCC TROT program (5T32CA160001), a GMTEC Postdoctoral Researcher Innovation Grant, and is an HHMI Hanna Gray Fellow.

---

**Extended Figure 1.**
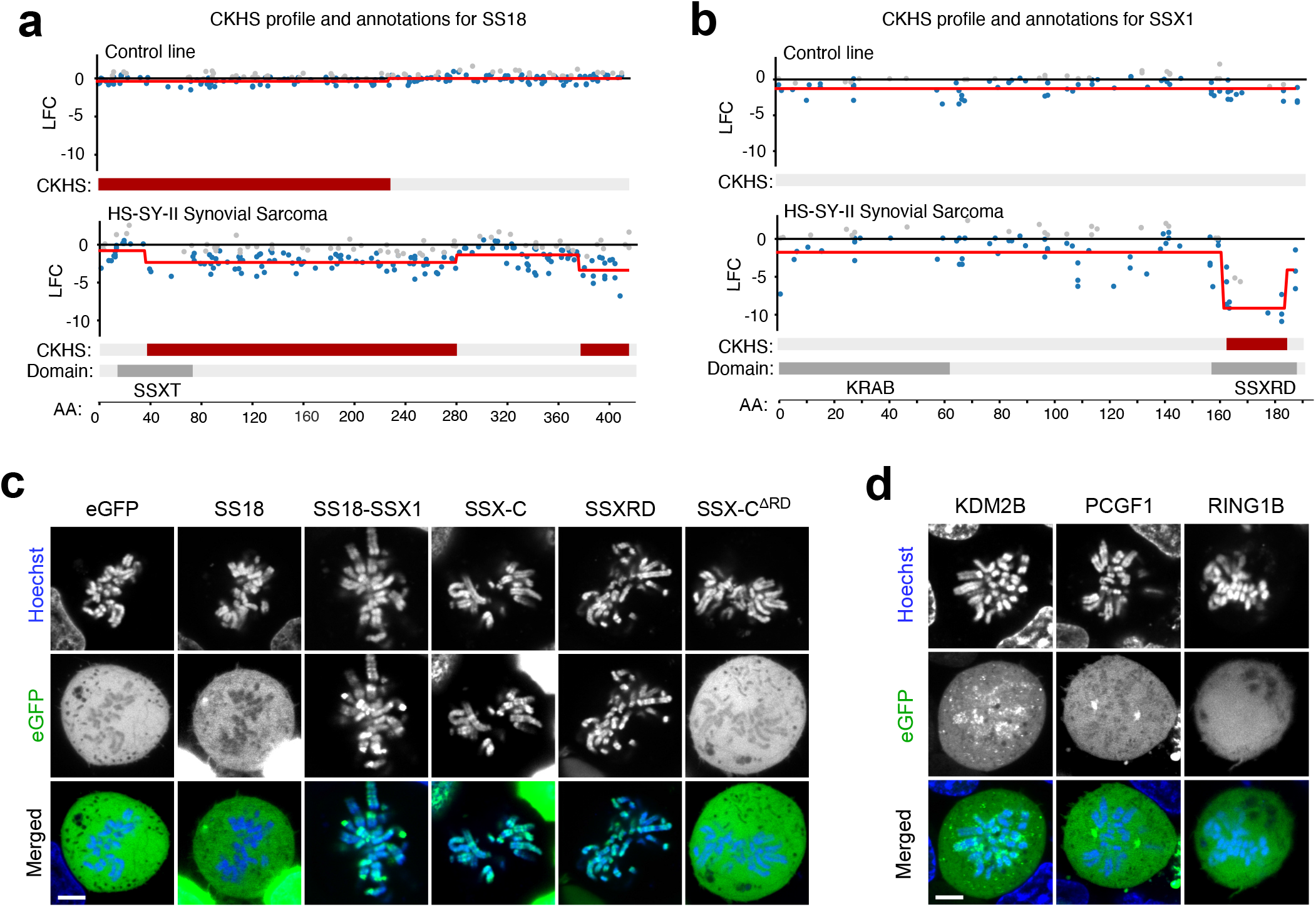
**a), b)** CRISPR knock-out hypersensitive (CKHS) regions and PFAM domain annotation for SS18 (a) and SSX1 (b) in control, fusion negative cell line (KHOS-240S, osteosarcoma cell line) and in HS-SY-II synovial sarcoma cell line harbouring an SS18-SSX1 fusion. CKHS regions are highlighted in dark red. **c), d)** Live confocal imaging images of metaphase HEK293T expressing eGFP fused constructs for SS18, SS18-SSX1, SSX-C, SSXRDand SSX-C^ΔRD^(c)orKDM2B, PCGF1 and RING1B (d). DNA is stained using Hoechst 33342. Scale bars correspond to 5μm.

**Extended Figure 2.**
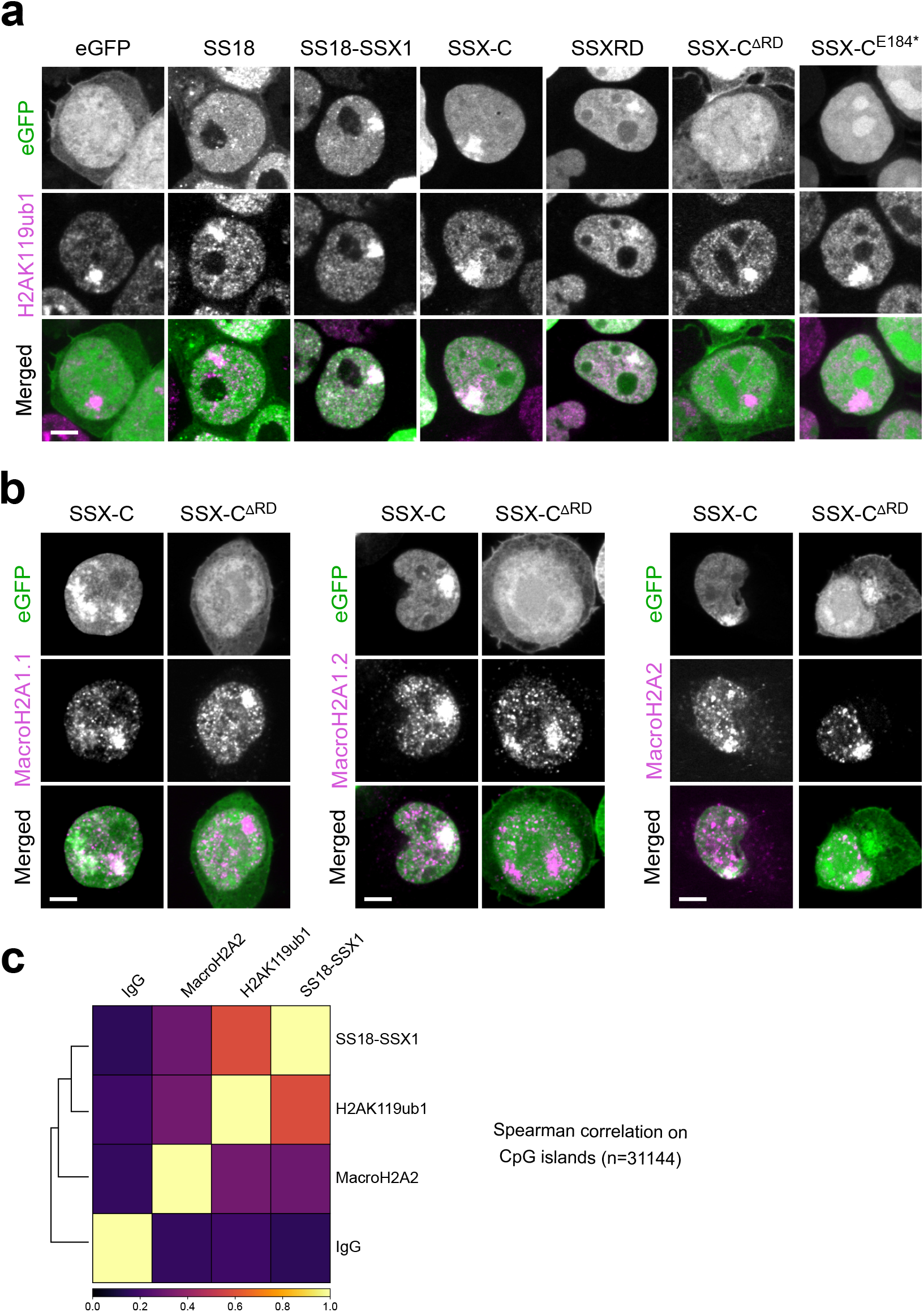
**a), b)** eGFP, H2AK119ub1 or histone MacroH2A immunofluorescence of HEK293T cells expressing the indicated eGFP constructs. Bottom panel displays merge channels with eGFP (green) and H2AK119ub1 (a) or histones MacroH2A (b) (magenta). Scale bars correspond to 5μm throughout the figures. **c)** Heatmap of Spearman correlation coefficients from bigWig coverages computed over genome CpG islands downloaded from UCSC genome track (https://genome.ucsc.edu/).

**Extended Figure 3.**
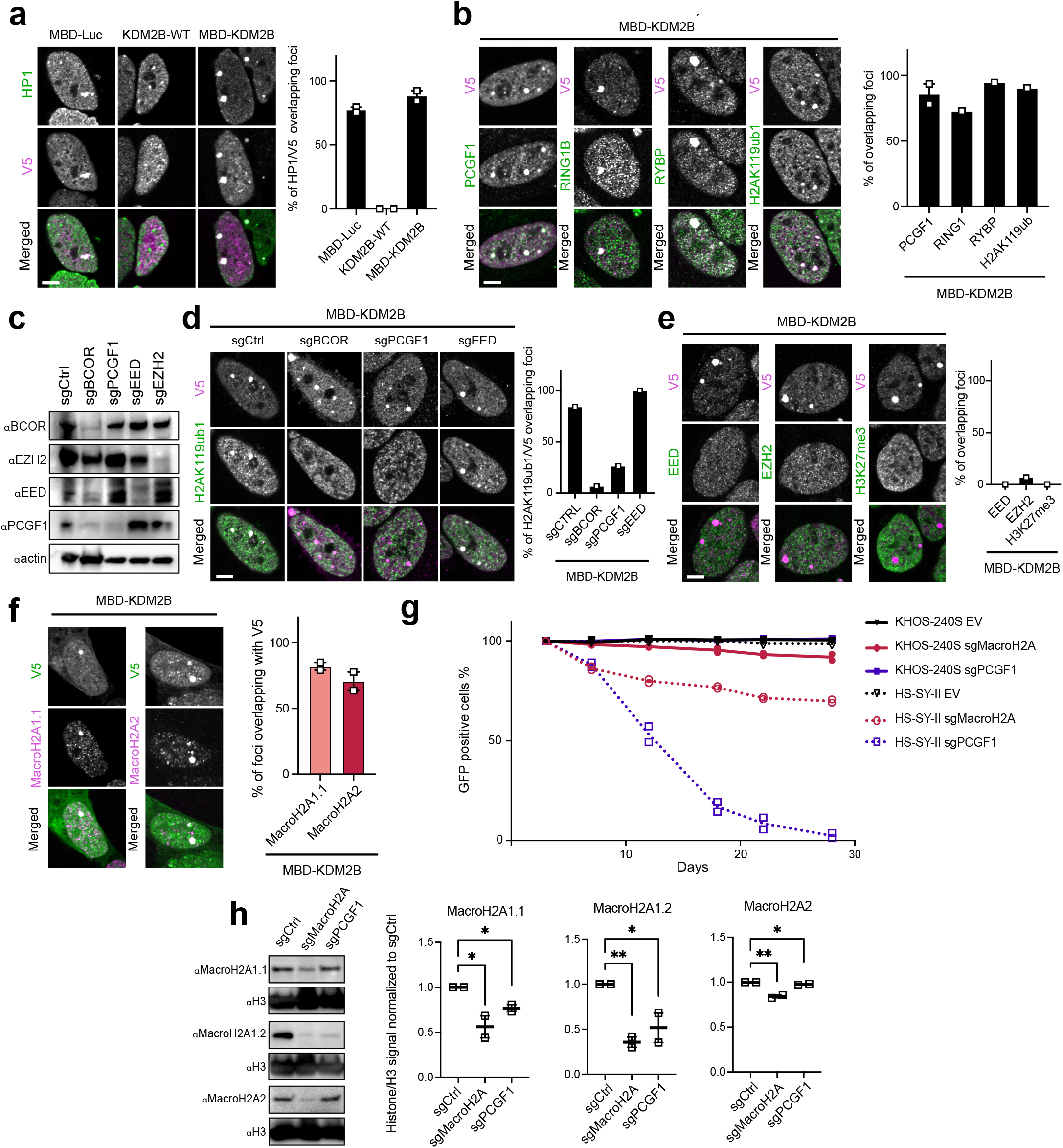
**a)** Left, Immunofluorescence for the MBD-Luc and KDM2B constructs or for KDM2B-WT fused to a V5 tag (V5, magenta) and HP1 (green). Scale bars represents 5μm throughout the figure. Right, quantification of the percentage of V5 foci overlapping HP1 foci in 2 biological replicates. Data represents the mean and the standard error of the mean (S.E.M). **b)** Left, Immunofluorescence of MBD-KDM2B (V5, magenta) with PCGF1, RING1B, RYBP and H2AK119ub1 (green). Right, quantification of the percentage of foci overlapping a V5 foci in 2 or 1 biological replicates. Data represents the mean and S.E.M when applicable. **c)** Western Blot of whole cell extracts from HS-SY-II-Cas9 cells expressing the various sgRNAs revealed using BOOR, EZH2, EED, PCGF1 or Beta-actin antibodies. **d)** Left, Immunofluorescence for MBD-KDM2B (V5, magenta) in the presence of different sgRNAs (resulting in eGFP background fluorescence) with H2AK119ub1 (green). Right, quantification of the percentage of H2AK119ub1 foci overlapping V5 foci in one biological replicate. **e)** Left, Immunofluorescence of MBD-KDM2B (V5, magenta) with EED, EZH2 or H3K27me3 (green). Right, quantification of the percentage of foci overlapping an V5 foci in 1 biological replicate. Data represents the mean. **f)** Left, Immunofluorescence of MBD-KDM2B (V5, magenta) with histones MacroH2A1.1 or MacroH2A2 (green). Right, quantification of the percentage of foci overlapping a V5 foci in 2 biological replicates. Data represents the mean and S.E.M. **g)** Cell competition assay performed in the osteosarcoma cell line KHOS-240S (fusion negative control) or in the synovial sarcoma line HS-SY-II transduced with an empty sgRNA as control or with guides targeting MacroH2A isoforms (sgMacroH2A) or PCGF1. **h)** Left, Western blot of histone acid extracts from HS-SY-II-Cas9 cells expressing sgCtrl, sgMacroH2A or sgPCGFI revealed with MacroH2A1.1, MacroH2A1.2, MacroH2A2 and H3 antibodies. Right, Signal quantifications using H3 as normalization, ratios were later further normalized to sgCtrl. Data represents the mean and the standard error of the mean (S.E.M) of two biological replicates. Asterisks represent p-values of unpaired one-tailed t-test between groups (* p< 0.05 and ** p< 0.01).

**Extended Figure 6.**
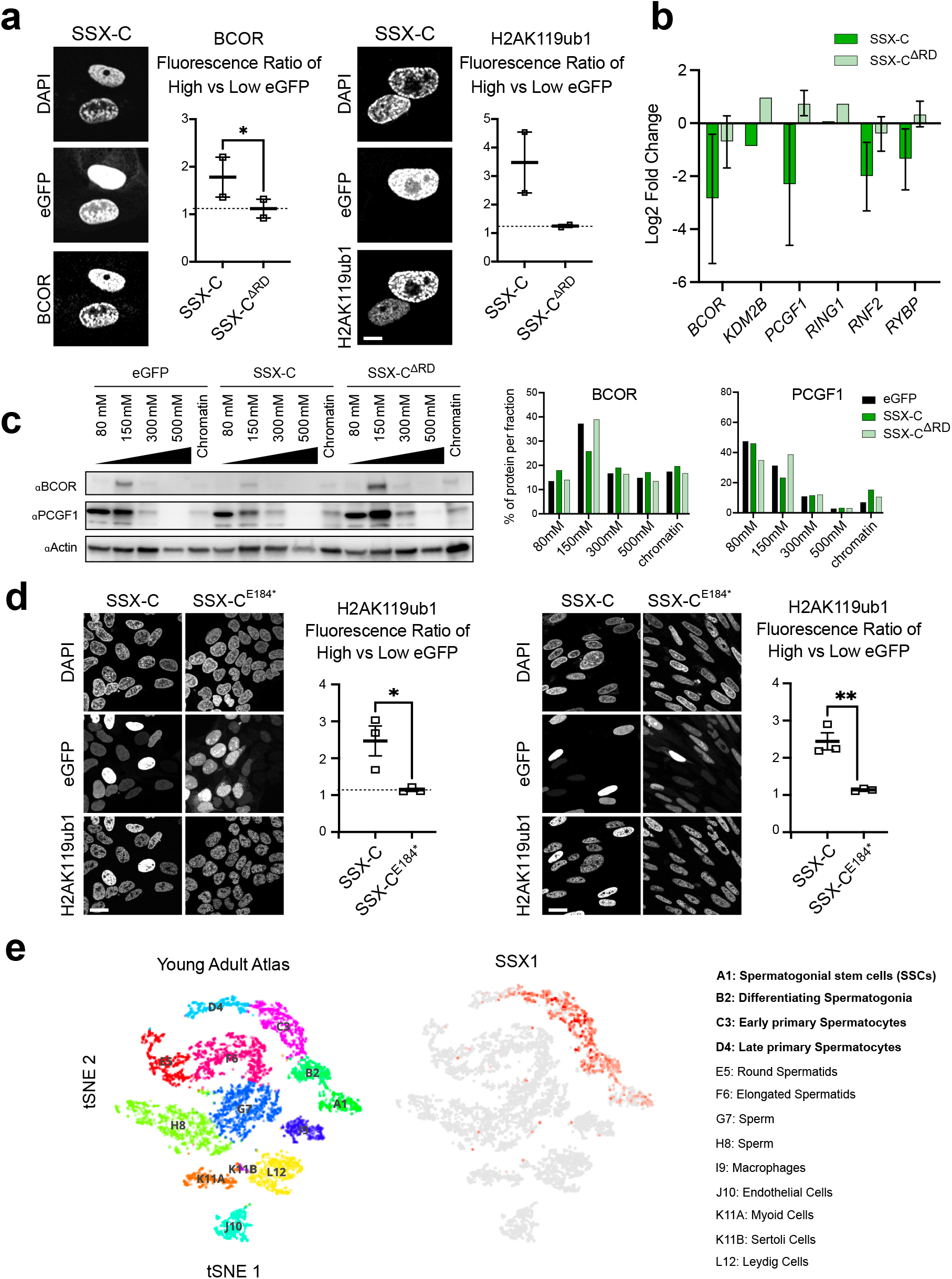
**a)** Left, Immunofluorescence against BCOR and H2AK119ub in mesenchymal stem cells (MSCs) expressing the indicated eGFP fused constructs with eGFP signals and nucleus stained with DAPI (grayscale). Right, quantification of BCOR or H2AK119ub1 fluorescence ratio in high versus low eGFP cells in 2 biological replicates. Data represents the mean and S.E.M. Asterisks represent p-values of paired one-tailed t-test between groups (* p< 0.05). Scale bars correspond to 10μm throughout the panel. **b)** qRT-PCR displaying Log2 fold change of mRNA levels normalized by *GAPDH* in MSCs expressing eGFP-fused constructs (eGFP, eGFP-SSX-C and eGFP-SSX-C^ΔRD^) relative to eGFP expressing cells in two biological replicates. Data represents the mean and the standard error of the mean (S.E.M). c) Salt extraction assay in HS-SY-II expressing eGFP, eGFP-SSX-C and eGFP-SSX-C^ΔRD^. Proteins were detected by western blot using with BCOR, PCGF1 or Beta-actin (loading control) antibodies. **c)** Quantification of the protein distribution in the various fractions of the salt extraction for BCOR or PCGF1. Data represents the percentage of total protein levels. **d)** Immunofluorescence against H2AK119ub in HS-SY-II (left) or SYO-I (right) cells expressing the indicated eGFP-fused constructs (eGFP-SSX-C and eGFP-SSX-C^E184^*). On the right of each panel of IF images are quantifications of the H2AK119ub1 fluorescence ratio in high versus low eGFP cells in 3 biological replicates. Data represents the mean and the standard error of the mean (S.E.M). Asterisks represent p-values of paired one-tailed t-test between groups (** p< 0.01) **e)** Left, tSNE and clustering analysis of combined single-cell transcriptome data from human testes (n = 6490) from (Guo et al., 2018). Each dot represents a single cell and is colored according to its cluster identity as indicated on the figure key. The 13 cluster identities were assigned based on marker gene expression. Right, SSX1 expression pattern projected on the tSNE plot. Red indicates high expression and gray indicates low or no expression.

**Extended Figure 7.**
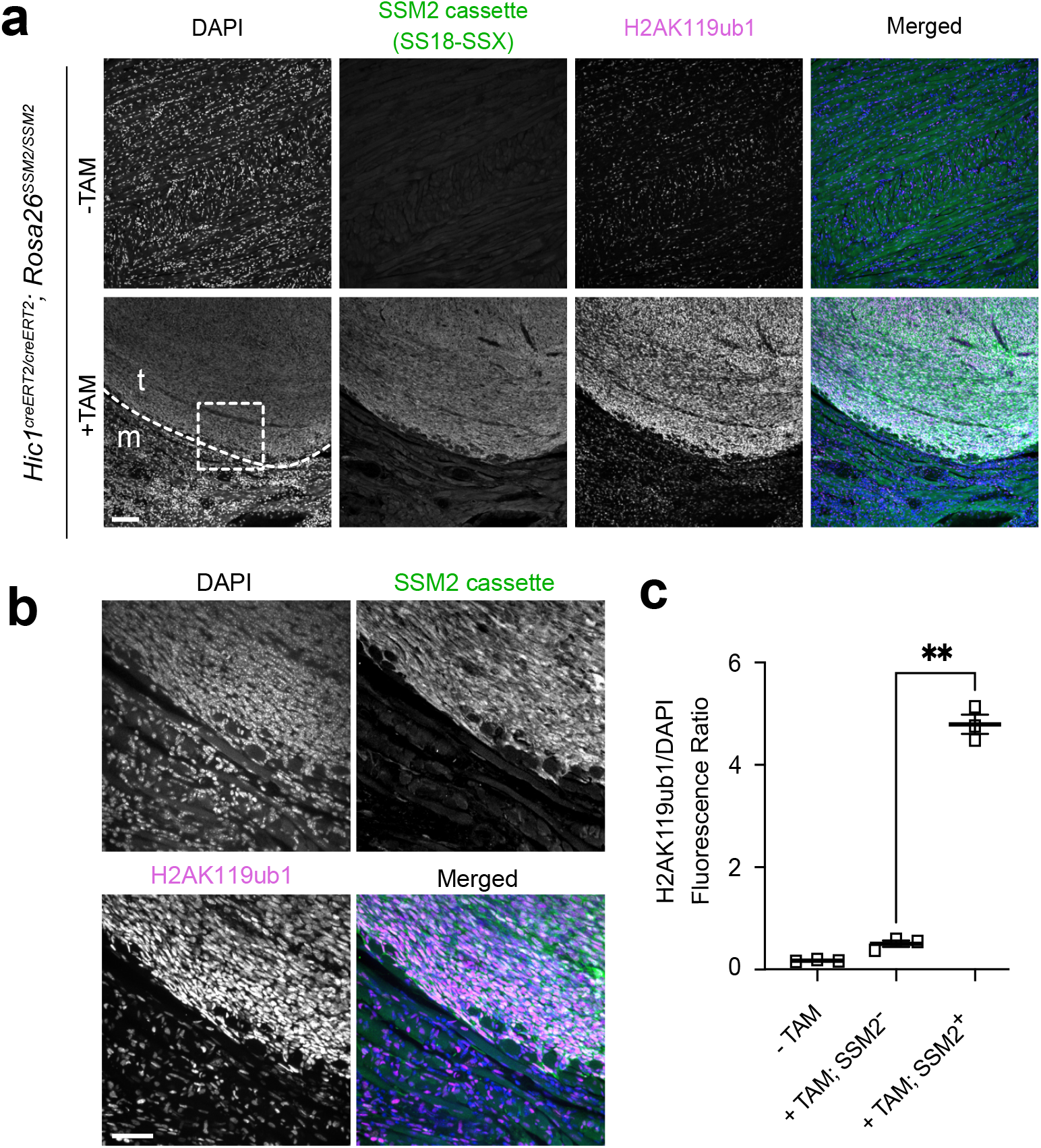
**a)** Immunofluorescence of *Hic1*^*creERT2/creERT2*^; *Rosa26*^*SSM2/SSM2*^ mice at 16-week endpoint tongue tissue showing left, samples from control mice not treated with tamoxifen (-TAM) (upper panel) or from tamoxifen treated (+TAM) mice expressing the SSM2 cassette (human SS18-SSX2) embedded in striated muscle (lower panel). The cells are stained for DAPI, SSM2 and H2AK119ub1. The scale bar represents 100 μm. **b)** Close-ups of images shown in the panel above, area corresponds to dashed line in (a). **c)** Quantification of H2AK119ub signal intensity normalised to DAPI signal intensity in 3 biological replicates in -TAM control mice, or +TAM tongue muscle or adjacent tumours (SSM2 negative or positive respectively). Asterisks represent p-values of paired one-tailed t-test between groups (** p< 0.01).

**Table S1.**
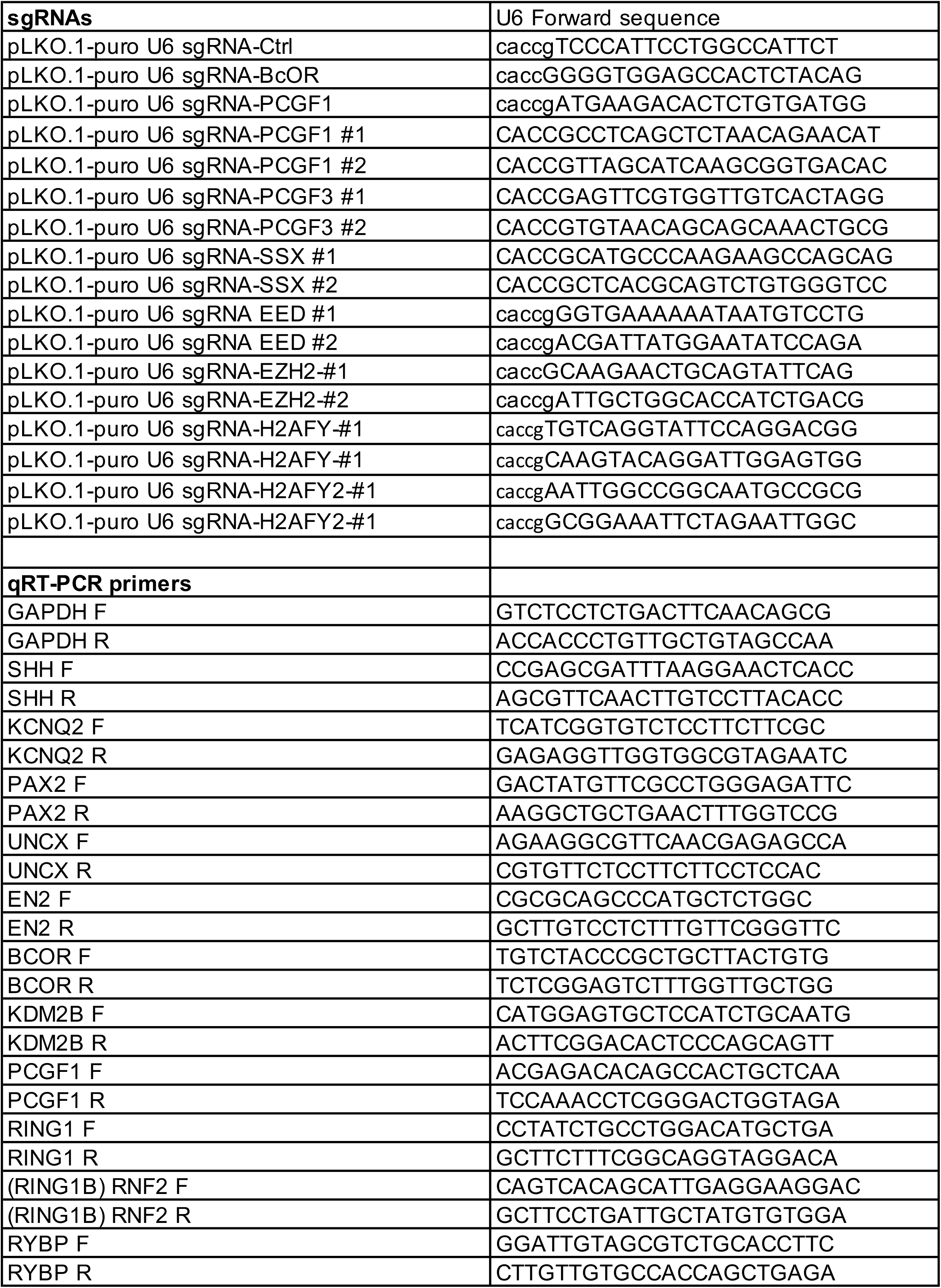
Sequence of DNA oligos used in this study.

**Table S2.**
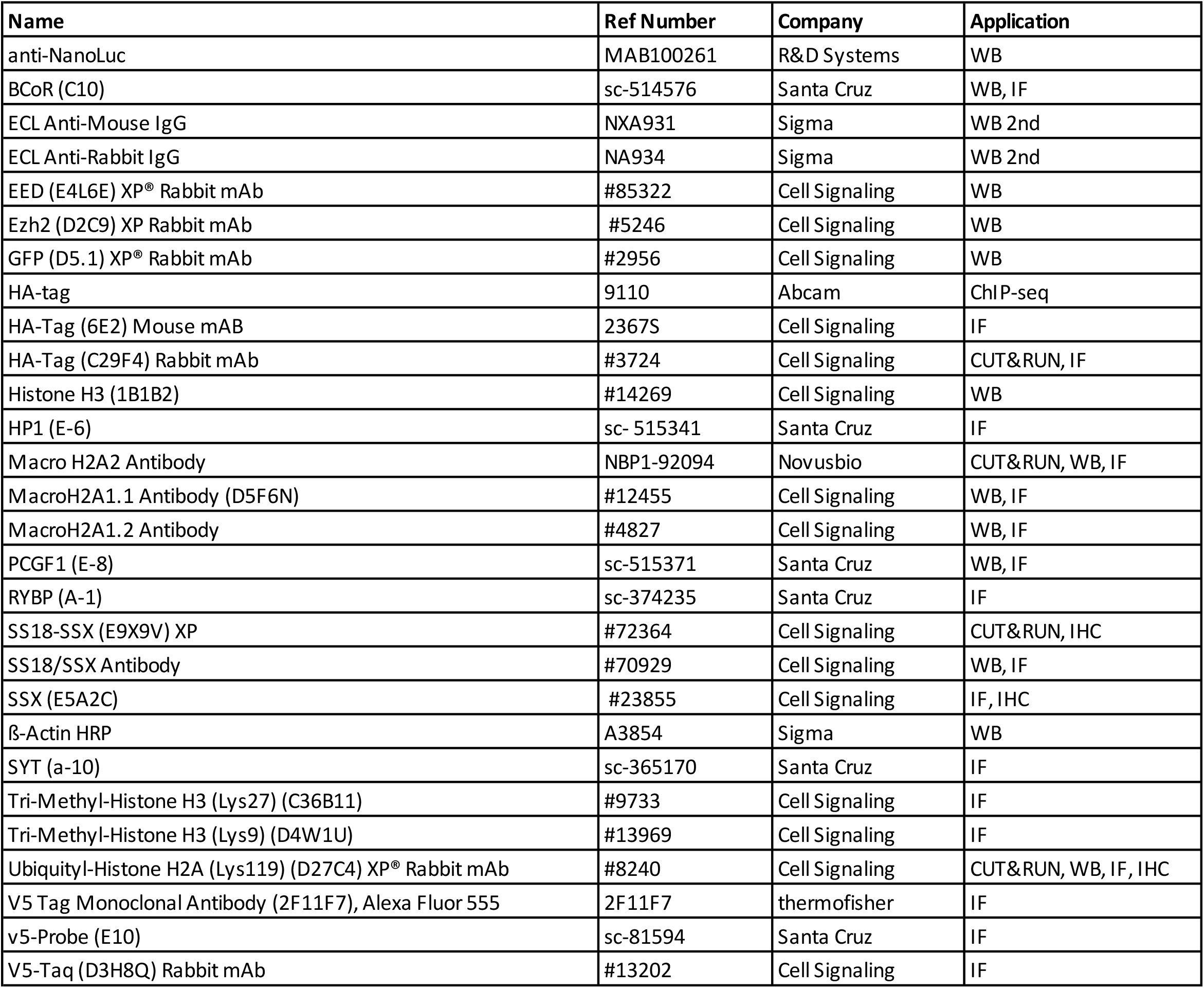
List of Antibodies used in this study.

